# Hydrogel–Electromagnetic Biohybrid Systems Direct Neural Morphogenesis across Central and Peripheral Systems

**DOI:** 10.1101/2024.09.29.615659

**Authors:** Bjarke Nørrehvedde Jensen, Tong Tong, Yun Tang, Grith Skovborg, Yuge Zhang, Kjeld Kaj Klompmaker, Cecillie Linneberg Matthiesen, Jens Toft Eschen, Kirstine Juul Elbæk, Yuting Wang, Lone Tjener Pallesen, Dror Seliktar, Di Sun, Mingdong Dong, Christian Bjerggaard Vægter, Anders Rosendal Korshøj, Menglin Chen

## Abstract

Biohybrid materials that integrate structural and bioelectronic cues across scales offer new routes to regulate neural development and repair. We present a hybrid, hydrogel–electromagnetic stimulation platform with a 3D hierarchically anisotropic structure, combining FLight bioprinted hydrogel microfilaments with a melt-electrowritten wireless electromagnetic stimulation system, to guide neural morphogenesis across central and peripheral systems at cellular and tissue scale. In the central nervous system, the soft microfilaments selectively promoted healthy, white-matter-like neurite extension rather than glioblastoma-derived outgrowth in human cortical brain tissues, while the electrically stimulation further enhanced the hippocampal neurosphere networking along the microfilaments. As a peripheral nerve conduit, the system directs dorsal root ganglion alignment in vitro and, in rat sciatic defects, drives a coordinated pro-regenerative response that improves axonal guidance, remyelination, and functional recovery. This multiscale biohybrid strategy demonstrates how synergistic integration of soft biomaterials and systematic, wireless bioelectric cues can non-invasively regulate neural morphogenesis, establishing new design principles for next-generation bioelectronic interfaces and regenerative medicine.

## Introduction

The extracellular matrixes (ECM) in both the central and peripheral nervous systems are responsible for maintaining structural integrity and homeostasis. They form the scaffolding for guiding cell migration, differentiation, and axonal outgrowth. Failure or defects in the brain ECM may result in diseases such as stroke, multiple sclerosis, Parkinson’s disease, or brain injuries [1–3]. Damage to the peripheral nerve ECM from injuries or diseases often lead to loss of function, even after undergoing autograft surgical treatment [4–9]. Proper reproduction of the native extracellular matrix is imperative to ensure high neural cell viability and correct phenotypic expression. As such, the structural, chemical, and mechanical properties of the native tissue must be recreated [1]. Furthermore, structural morphology greatly influences cellular morphology [10, 11]. Studies have shown that, depending on the cell type, structural features as small as 33 nm and as large as 100 µm can influence cellular morphology [12, 13]. As such, nano- and microgrooves in surfaces and nano/microfibers are utilized in neural tissue engineering to align neurons and to mimic ECM structures that occur in mature nervous systems. Such anisotropic structures have been fabricated using various techniques such as electrospinning [14, 15], melt electrowriting [16–18], freeze drying [19, 20], electric fields [21], and bioprinting [22, 23]. Due to the difficulty in structuring hydrogels with <1 kPa modulus, limited work has been done on defined *in vitro* patterns with physiologically relevant moduli [24–28]. Consequently, the structures remain simple and do not resemble physiological structures [29]. Biofabrication utilizing light filamentation has emerged as a promising technique for rapid fabrication of aligned hydrogel structures. Liu *et al.* demonstrated this by fabricating highly anisotropic and aligned hydrogel microfilaments using light-based projection. The microfilaments, with diameters matching cell sizes (2-30 µm), exhibit highly unidirectional structural characteristics [32]. These structures have been demonstrated on muscle tissue, tendons, and for advanced versions of nerve guide conduits [30–33].

The electrical activity of cells plays a vital role in neural proliferation, migration, repair, and differentiation at different stages [34]. Studies show that electrical stimulation during tissue development and regeneration can activate neural regenerative pathways resulting in cell outgrowth and migration, axonal directionality, and extension, promote neurotrophic factor production, increase neural plasticity, and cause an upregulation of regeneration-related genes [35, 36]. Delivery of direct electrical stimulation can be disruptive to the damaged tissue, can cause further damage, and has the potential risk of inducing infection both during stimulation and during the retrieval of the stimulation device [37–40]. As such wireless stimulation methods are a growing field of research that aims to solve the limitations of direct stimulation [38]. One of the promising techniques utilizes electromagnetic (EM) induction to induce stimulating currents in materials within the damaged site [41]. EM induction has been used to create fully implantable and bioresorbable neural stimulators. A magnesium coil was coated with wolfram and encapsulated in PLGA to isolate it for implantation. Megahertz frequency magnetic fields were converted to direct current-like currents through up to 15 cm of native tissue. The coil was connected to a nerve cuff for transient stimulation of the sciatic nerve of mice. Wireless electrical stimulation (monophasic, 200 µs pulse, 20 Hz frequency) was delivered to an injured nerve for 1 h per day for 1-, 3-, and 6-days post-injury. The results showed significantly increased rates and degrees of nerve regeneration in addition to increased recovery of muscle function [42–45]. Further utilizing of an anisotropic coil design, wireless EM stimulation yielded 73.2% increased PC12 neural outgrowth over 7 days of stimulation [16]. However, little focus is put into the 3D morphology of the inductive materials and as such does not utilize the combined effect of topographical guidance with electrical stimulation.

The convergence of different 3D printing technologies has great potential to exploit multi-function of hybrid materials in hierarchical structure. In this study, soft, anisotropic hydrogel microfilaments were integrated with wireless EM stimulation, for the first time, to guide neural morphogenesis across central and peripheral nervous systems at both cellular and tissue scale (**Figure 1**). Gelatin methacrylate (GelMA) microfilaments’ swelling ratio, mechanical modulus were first characterized. The anisotropic EM stimulation coil, fabricated by melt electrowritting (MEW) polycaprolactone (PCL) followed with coating of 80nm gold (Au), was then integrated in filamented light (FLight) bioprinting of GelMA microfilaments and their biocompatibility and ability to guide 3D cell alignment under EM stimulation were assessed. Further investigations of *ex vivo* white matter axonal guidance in human brain tissue, *in vitro* hippocampal neurosphere networking, and a peripheral neural guide conduit were conducted. To translate our *in vitro* findings into a therapeutic context, this integrated system was investigated in a critical-sized rat sciatic nerve defect. We systematically assessed the regenerative efficacy by examining key outcomes that ranged from the quality of structural and cellular repair to the recovery of physiological nerve conduction, and finally, the functional reinnervation of downstream muscle.

**Figure 1.**
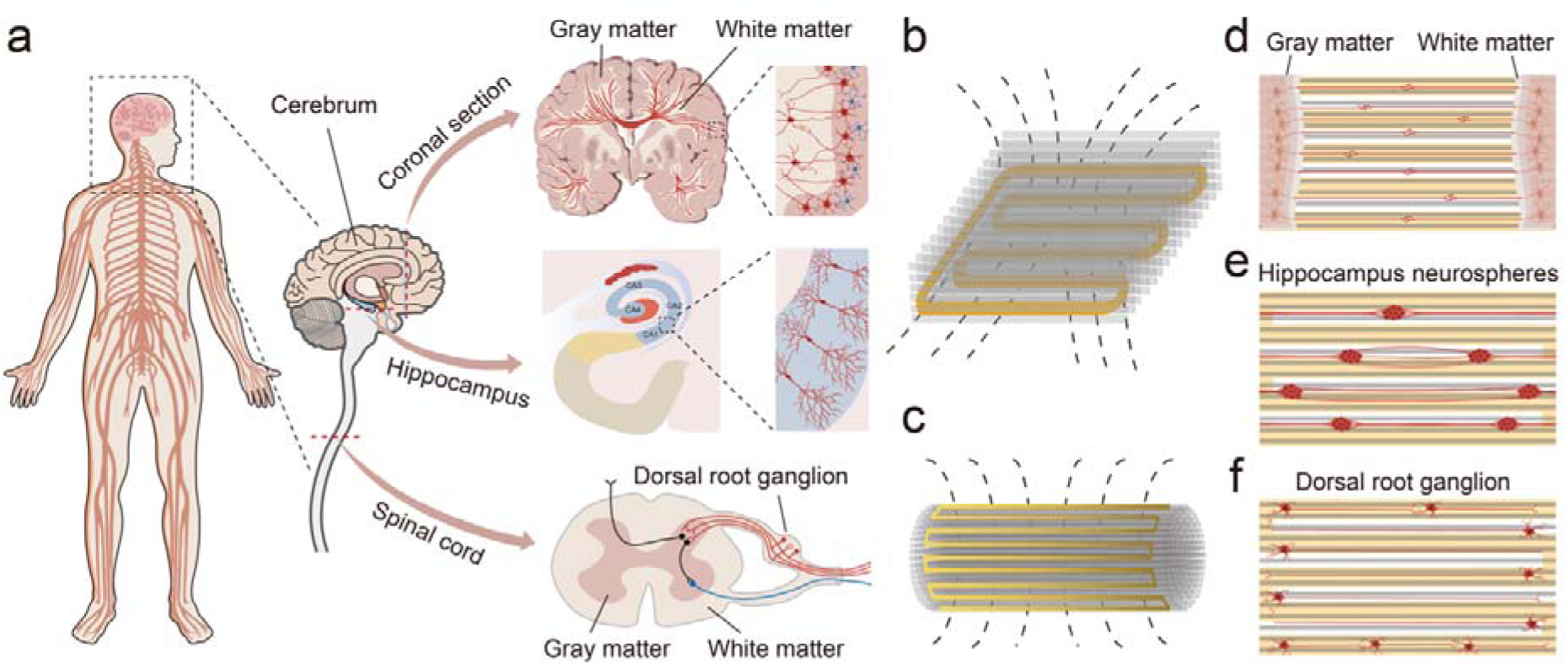
Oriented hydrogels with wireless electromagnetic (EM) stimulation for guided neural morphogenesis across central and peripheral nervous systems (**a**). Anisotropic hydrogel filaments integrated with EM stimulation coils (**b, c**) for topographically guided human brain tissue white matter connectivity (**d**), hippocampus neurosphere formation (**e**), and dorsal root ganglion neural outgrowth (**f**).

## Results and discussion

### MEW/FLight Fabrication, Mechanical Characterization and Biocompatibility

MEW, which allows the creation of highly organized fiber architectures having micron-scale resolution, has been used to confer outstanding mechanical properties to low-stiffness hydrogels. Only until recently, MEW has been combined with light based bioprinting, such as volumetric bioprinting, a rapid light-based bioprinting technology to sculpt hydrogel into free-from 3D structures.[46] Here, we combine MEW with another light based bioprinting, FLight biofabrication technique [33], to integrate hybrid materials into a 3D hierarchically anisotropic structure for topological/bioelectrical dual-functional cellular guidance. (**Fig 2**) First, the PCL anisotropic coil was fabricated using MEW (**Fig 2A,i**) creating fibers 30 µm in diameter of PCL in a 40-layer stack (as walls) of fibers with a closed loop configuration enables EM induction of current in the coil upon application of an alternating magnetic field. The coil was rendered conductive through glancing angel deposition (GLAD) coating with 80 nm of Au, confirmed by Scanning Electron Microscopy-Energy Dispersive X-ray Spectroscopy (SEM-EDX).(**Fig 2B**) Then, the MEW printed EM coil, either as a patch or rolled up in a tube mold filled with GelMA solution, was placed on a home-made PDMS holder in a bioprinting cuvette and cooled down to 4 °C. (**Fig 2A,ii**) Last, the printing cuvette was placed into the printer, with the long walls of EM coil align with the beam projection (**Fig 2A,iii, 2C**). Finally, after FLight bioprinting, well oriented GelMA microfilaments (GelMA-F) were created along the long walls of the MEW printed EM coil (**Fig 2D**). The voltage output of the EM stimulation system with hydrogel encapsulation shows a decrease (folds) in the induced voltage upon GelMA encapsulation (**Figure 2D**), where 7.2 mV is induced at 90 kHz, 10V. The GelMA in phosphate buffered saline (PBS, conductivity close to that of common cellular media and body fluid [47, 48]) functions as a conductive electrolyte allowing for ionic currents through the system.

**Figure 2.**
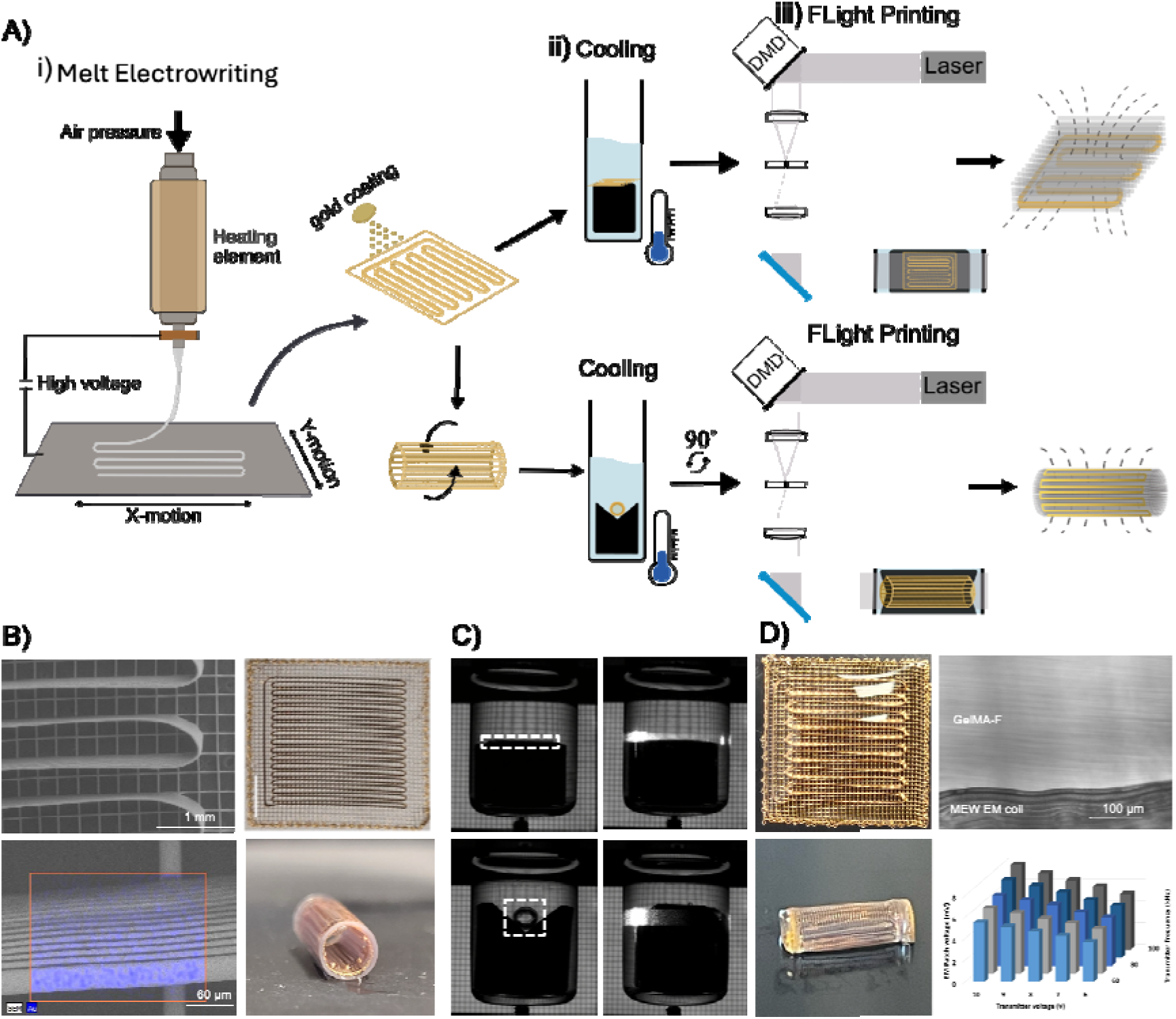
Convergence of MEW and FLight processes to fabricate GelMA-F/EM ptach or conduit. **A**) Graphical overview of: i) the fabrication of melt electrowritten (MEW) coils, Au coating, rolling and ii) their subsequent incorporation into the FLight process by first placing the MEW coils in GelMA solution on a home-made PDMS holder (black)in a printing cuvette, cooled to 4 °C to fully gelled and iii) placed in the printer with the long walls of MEW align with the laser projection. The same process can be applied in the presence of cells. **B**) SEM/EDX images and photographs of MEW coil as a patch or rolled up and inserted in a tube mold. **C**) Digital images of the printing vials containing GelMA and a MEW coil patch/tube when inserted into the printer and when light projections are initiated (the tube was turned 90 °) from the left side. **D**) Photographs of the patch or tube (mold removed) after FLight printing, and a light microscopy image of printed GelMA-F along with the long walls of MEW coil, together with the voltage output of when encapsulated in GelMA-F/EM patch.

The obtained GelMA-F bundles exhibit an exceptionally high degree of fiber alignment compared to the isotropic structure of bulk GelMA (GelMA-B), as shown in the brightfield images and Fast Fourier Transform (FFT) analysis inserts **(Figure 3.a-d)**.

**Figure 3.**
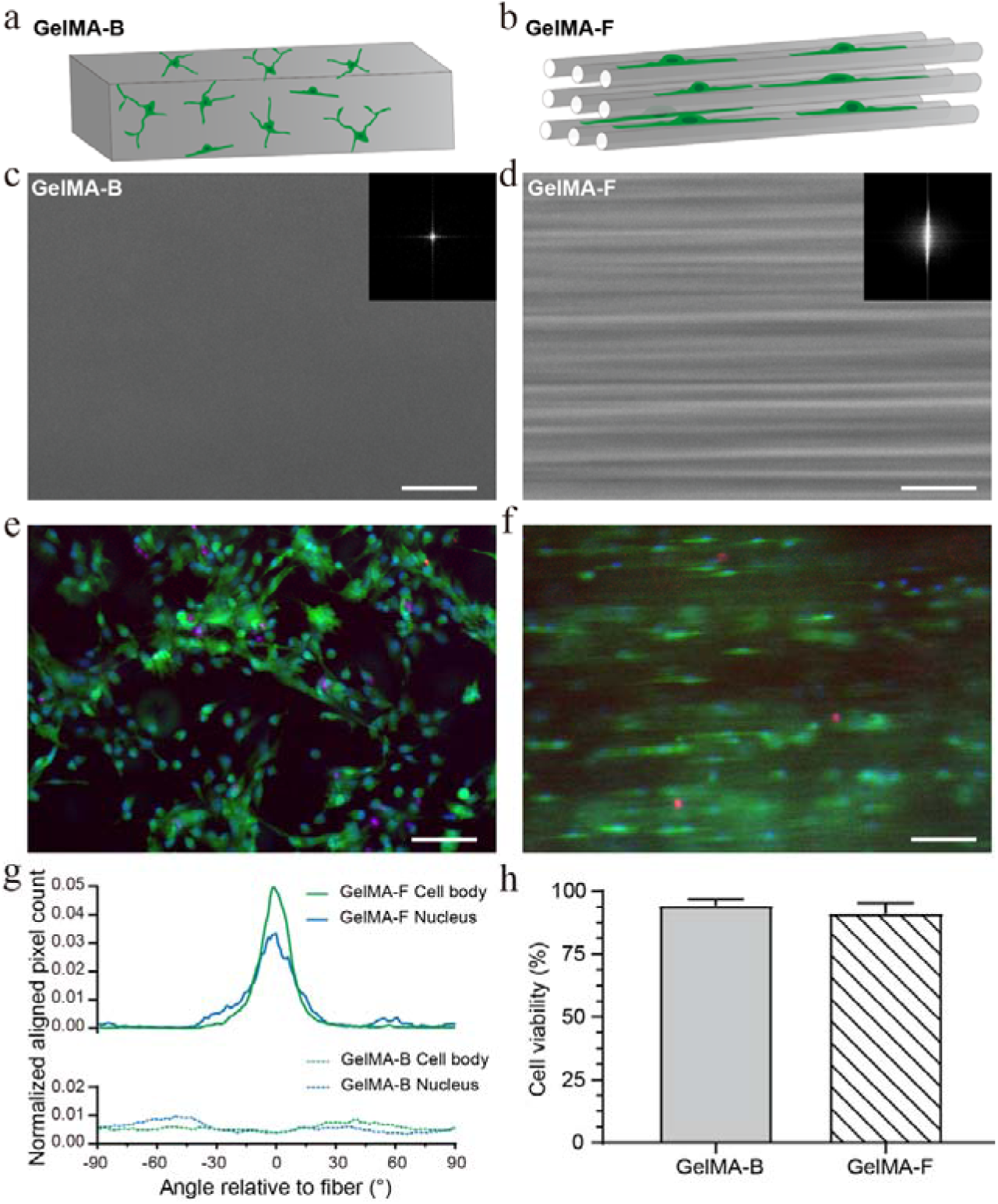
Biocompatibility and topographical guidance of filamented GelMA-F. **a-b)** Schematic of eGFP-MSC cells in GelMA-B and GelMA-F. **c-d)** Brightfield images with FFT as insets of 4% w/v GelMA-B and GelMA-F. **e-f)** Fluorescence images of eGFP-hMSC cells in GelMA-B and GelMA-F after 6 days of culture. red = propidium iodide (dead cells). **g)** Cell alignment quantification after 6 days of culture. **h)** Cell viability based on live/dead imaging after 6 days of culture. Scalebars = 100 µm.

The swelling ratio of GelMA-B and GelMA-F hydrogels at 2%, 3%, and 4% w/v were investigated (**Figure S1.a-b**). Minimal differences in swelling ratio between 2%-4% w/v GelMA are observed, while the increased swelling of GelMA-F may be attributed to the interstitial space created in GelMA-F hydrogels [33] which accommodated larger amounts of water between the loosely bound fibers similar to how granulated hydrogels swell passing the polymer matrix swelling capacity [49]. Young’s modulus of GelMA was found to increase (range from 1 to 20 kPa), along the concentration increase from 2% to 4% w/v. (**Figure S1.a-b**).

Sculpting the 3D microenvironment to mimic the native tissue structure enhance cellular processes and function [10]. Connective tissues, fundamental in the body, possess hierarchical, anisotropic features reminiscent of fibers, along which cells grow and differentiate [11, 50]. Green fluorescent cells (eGFP-hMSCs) allowed for live-imaging were encapsulated in the GelMA-F and GelMA-B (**Figure 3.e-f**) to monitor the effect of topography upon the cell morphology. After 6 days, the fluorescent images show cells growing and spreading along the GelMA-F fibers, compared to the isotropic growth pattern in GelMA-B. (**Figure 3.g**). The fabrication and presence of the filamented hydrogel, GelMA-F, had no adverse effect on the viability of encapsulated cells (**Figure 3.h)**.

### Connectivity in the white matter of human cortical brain tissue

Anisotropic features are found throughout the white matter of the brain [51]. The main function of the white matter is to connect neurons from different brain regions to each other by myelinated axons. The white color comes from the abundant presence of lipids in the myelin sheaths made by oligodendrocytes [52]. The highly ordered axonal structure underlies white matter functionality and plays a major role in brain connectivity [53].

Human brain tissues from the cortex region infiltrated with glioblastoma cancer were placed on GelMA-F to investigate the outgrowth guidance. A concentration of 2% w/v GelMA-F was chosen as its Young’s Modulus of 1-3.5 kPa is close to that of white matter in native human cortical tissue (**Figure S1.c-d**) [54, 55]. A slice of brain tissue was cut into two pieces which were placed onto the GelMA-F, held by PTFE membrane culture insert, with a gap of 1-1.5 mm (**Figure 4.a,b**). The two half-slices were placed on top of GelMA-F with gray/white matter orientation perpendicular to the fiber direction, investigating the role of GelMA-F on the topographical guidance of the axonal pathways transitioning from the gray matter into the white matter. After 21 days of culture, tissue outgrowth on the PTFE membrane culture insert (**Figure 4.c,e**) or on GelMA-F (**Figure 4.d,f**) was observed. Larger holes in the coverage between the two brain tissue pieces are observed on the membrane inserts, while aligned and guided axonal connectivity is observed on GelMA-F.

**Figure 4.**
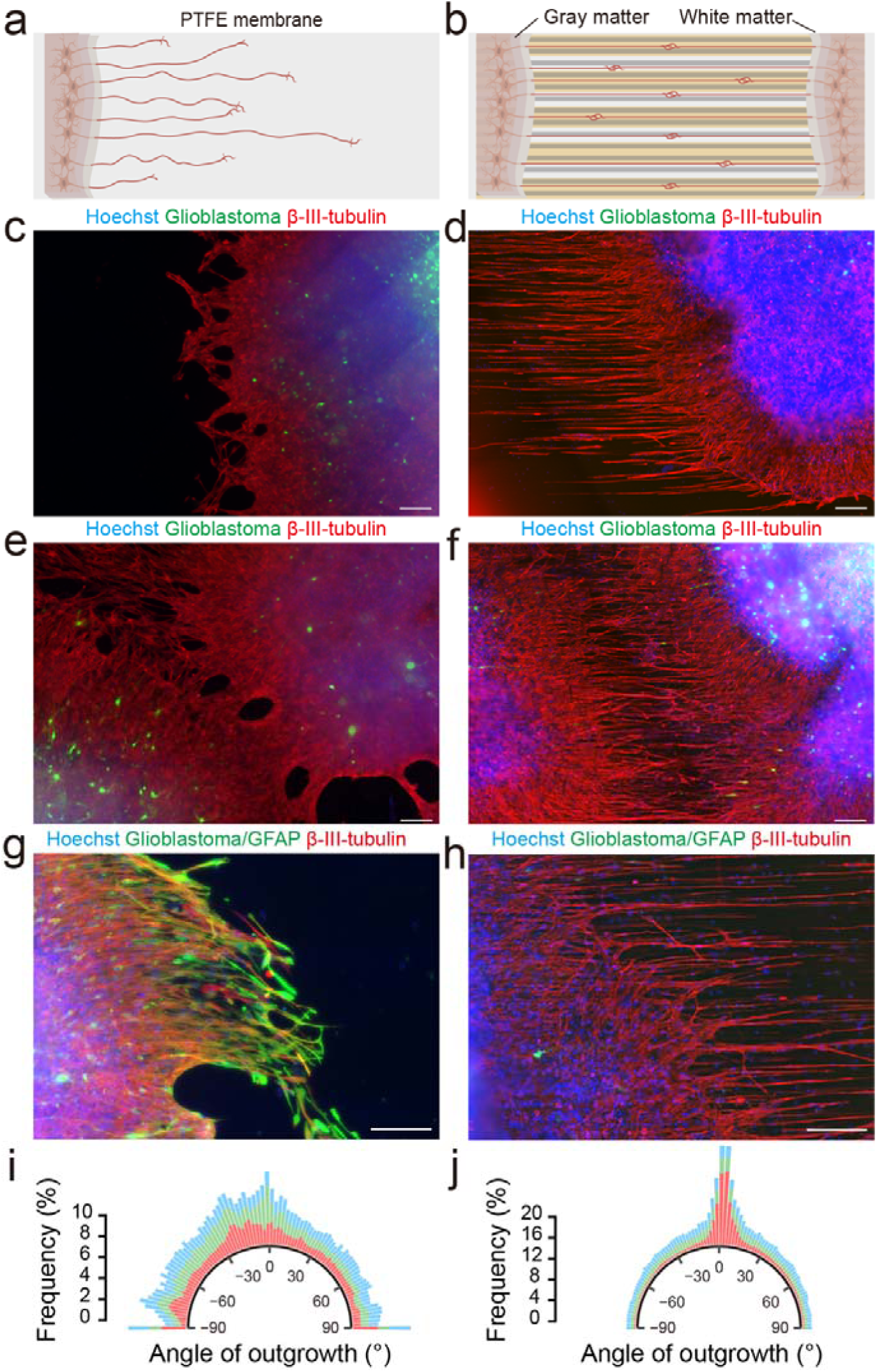
Guided human cortex brain tissue outgrowth on GelMA-F. **a-b)** Schematic of human brain cortex tissue cultured on PTFE membrane culture insert or GelMA-F. Outgrowth from slices of human cortex brain tissue after 21 days on membrane well-insert (**c** – side, **e** - middle), and on 2% w/v GelMA-F (**d** – side, **f** - middle). Blue = Hoechst, red = β-III-tubulin, green = glioblastoma. Outgrowth from slices of human cortex brain tissue after 21 days post-ICC-stained with GFAP on **g)** membrane well-insert, and **h)** 2% w/v GelMA-F. Blue = Hoechst, red = β-III-tubulin, green = glioblastoma/GFAP. Scalebar = 200 µm. **i-j)** Directionality analysis of the outgrowth from human cortex brain tissue.

We further stained with Glial Fibrillary Acidic Protein (GFAP), a marker of astrogliosis, following CNS injuries [56] and neurodegeneration, such as in Alzheimer’s disease [57, 58], and malignancies of glial origin, glioblastoma [59]. GFAP is also a key intermediate filament protein of glial cells involved in cytoskeleton remodeling and linked to tumor progression. Remarkably, very limited presence of GFAP was found on GelMA-F outgrowth but GFAP-positive glial cell outgrowth on PTFE membrane (stiffness in the MPa range) well-inserts (**Figure S2 and Figure 4.g-h**), suggesting GelMA-F selectively induced healthy neurite outgrowth rather than glioblastoma microtubes associated with tumor progression. A similar stiffness dependent glioblastoma expansion has been reported before [60], where glioblastoma has been found to preferentially expand in mechanically stiffer environments.

A clear alignment of the outgrowth with the GelMA-F fibers is observed, while isotropic outgrowth is seen on the PTFE membrane inserts. (**Figure 4.i,j**) Furthermore, the outgrowth is not directed when cultured on GelMA-B (**Figure S3**), confirming that the topographical guidance of the aligned GelMA-F fibers.

The human cortex brain tissue was then placed upon a GelMA-F encapsulated EM patch made of Au-coated MEW PCL coils (**Figure S4a,b**) to investigate the wireless EM stimulation on the outgrowth of brain tissue. After 7 days EM stimulation (7.2 mV, 90 kHz, 2 h/day, 7 days), tissue integration was seen onto the wall of the EM stimulation patch (**Figure S4c,d**), where both the nucleus and cell body appear to follow the direction of the anisotropic topographical features of the EM coil.

Therefore, GelMA-F guided neural outgrowth of human brain cortical tissue, demonstrating their potential use for expanding our understanding on how glial cells interact with axonal growth throughout neurodevelopment and investigating growth patterns of axonal connectivity in white matter of healthy and cancerous human brain tissue for physiological/pathological modeling.

Strikingly, GelMA-F appeared non-permissive for GFAP positive glial microtube outgrowth. The GelMA fibers show characteristics of radial glial fibers which are known to act as a scaffold for postnatal ventricular-subventricular zone (V-SVZ)-derived neuroblasts that migrate to the lesion site to promote neuronal regeneration. White matter maturation is a complex process combining changes in axonal diameter and packing, myelin thickness, and composition. The role of gray matter in modulating cortical activity through topographically guided white matter is not investigated in our study.

### Hippocampal neurosphere network

As an extension of the cerebral cortex situated deep into the temporal lobe, the hippocampus is vital for learning, memory, and spatial navigation. It is vulnerable to a variety of stimuli and hence is important clinically both diagnostically and therapeutically for cognitive decline and diagnosis of Alzheimer’s disease. Intriguingly, the hippocampus is one of the unique regions in brain where the neurogenesis continues even in adult life [61]. Microfluidic systems or microfabricated surfaces are widely used to guide neuronal connections to investigate basic neuronal circuits and functions [62–65]. However, while many of these systems are successful in generating circuits, they often lack the potential influence of the mechanical and molecular properties of the surrounding ECM. Here we investigate if GelMA-F can be used as a step towards a platform for generating guided hippocampal neurospheres by utilizing the aligned topography of GelMA-F (**Figure 5.a**).

**Figure 5.**
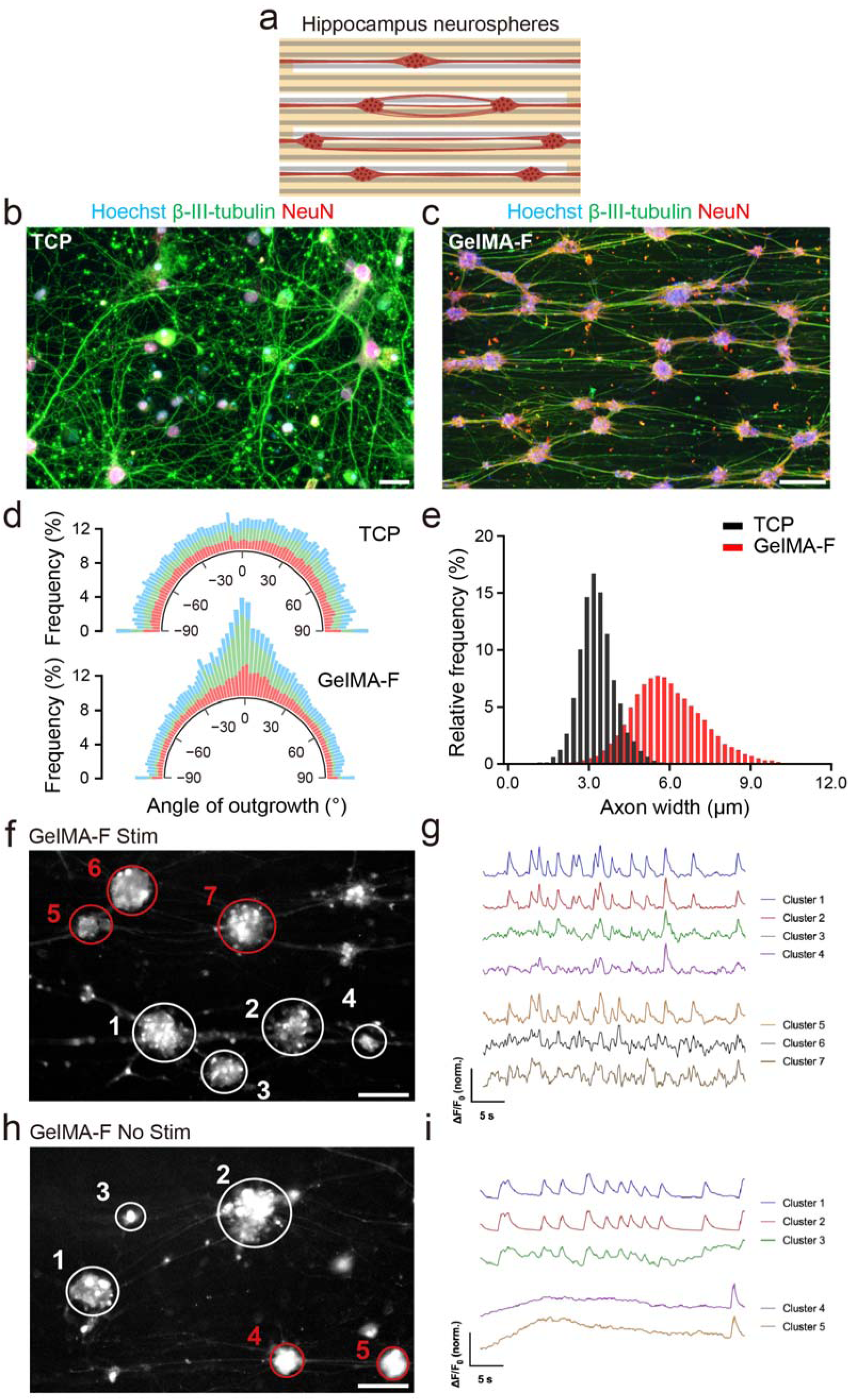
Hippocampal neuron neurosphere formation on GelMA-F. **a)** Schematic of hippocampal neurosphere network on GelMA-F. Confocal images of hippocampal neurons on TCP **(b)** and GelMA-F **(c)** after 14 days in culture. Scale bar, 25 µm (b) and 200 µm (c). **d)** Cluster axon angle distributions on TCP and GelMA-F. **e)** Cluster size distribution of neurospheres formed on 4% w/v GelMA-F and TCP. **g)** Representative synchronized spike trains of hippocampal neurospheres on GelMA-F after 14 days with EM stimulation. Scale bar 100 µm **h)** Representative synchronized spike trains of hippocampal neurospheres on GelMA-F after 14 days without stimulation. Scale bar 100 µm.

Hippocampal neurons were seeded onto laminin-coated tissue culture plastic (TCP) and 4% w/v GelMA-F with young’s modulus of 15-20 kPa (**Figure S1.c-d**), in the comparable range of previously reported rodent brains[66]. On TCP, hippocampal neurons appeared as singular cells with neurites spreading radially outward (**Figure 5.b**). The neurites, rather than spreading radially outwards from the neurospheres, were observed to form axon bundles aligned along GelMA-F (**Figure 5.c**). The cell and neurite directionality in relation to the direction of GelMA-F fibers was analyzed (**Figure 5.d**), where a clear directionality was found on GelMA-F compared to TCP, forming an aligned axon network among neurospheres. The axon width is further increased on GelMA-F as the axon bundle together between the neurospheres (**Figure 5.e**).

Further with EM stimulation (7.2 mV, 90 kHz, 2 h/day, 14 days), calcium imaging of hippocampal neurospheres cultured on GelMA-F enabled synchronous, faster spiking between sequential and adjacent neurospheres compared to non-stimulated group (**Figure 5.f-i**), confirming connectivity between sequential and adjacent neurospheres and preferential transmission along the fibers.

The hybrid system provides a straightforward neuromodulation platform for CNS neurosphere networks on a topographically controlled soft hydrogel surface, a feature that is uncommon in currently available systems [67–69], enabling facile establishment of neurospheres network as *in vitro* brain models to study cognitive diseases diagnostically and therapeutically.

### Neural guide conduit for peripheral neuron repair

It is well known that neural outgrowth is guided along anisotropic structures in peripheral neural regeneration [11]. GelMA-F was printed inside the EM coil [16] for guiding and stimulating peripheral neural regeneration (**Figure 6.a**). On TCP and unstructured GelMA-B surfaces, the dorsal root ganglion (DRG) neurons formed radially outward spreading neurites as commonly observed on isotropic surfaces [70–73]. While on the uniformly aligned GelMA-F surfaces, the neurites clearly aligned with the direction of the fibers (**Figure 5.b,c**). Importantly, without laminin or other basement membrane protein coatings traditionally required for the culture of DRG neurons on TCP surfaces, GelMA provides sufficient adhesion for DRG neurons for outgrowth reaching >3.2 mm over 7 days on GelMA-F (**Figure 6.d**).

**Figure 6.**
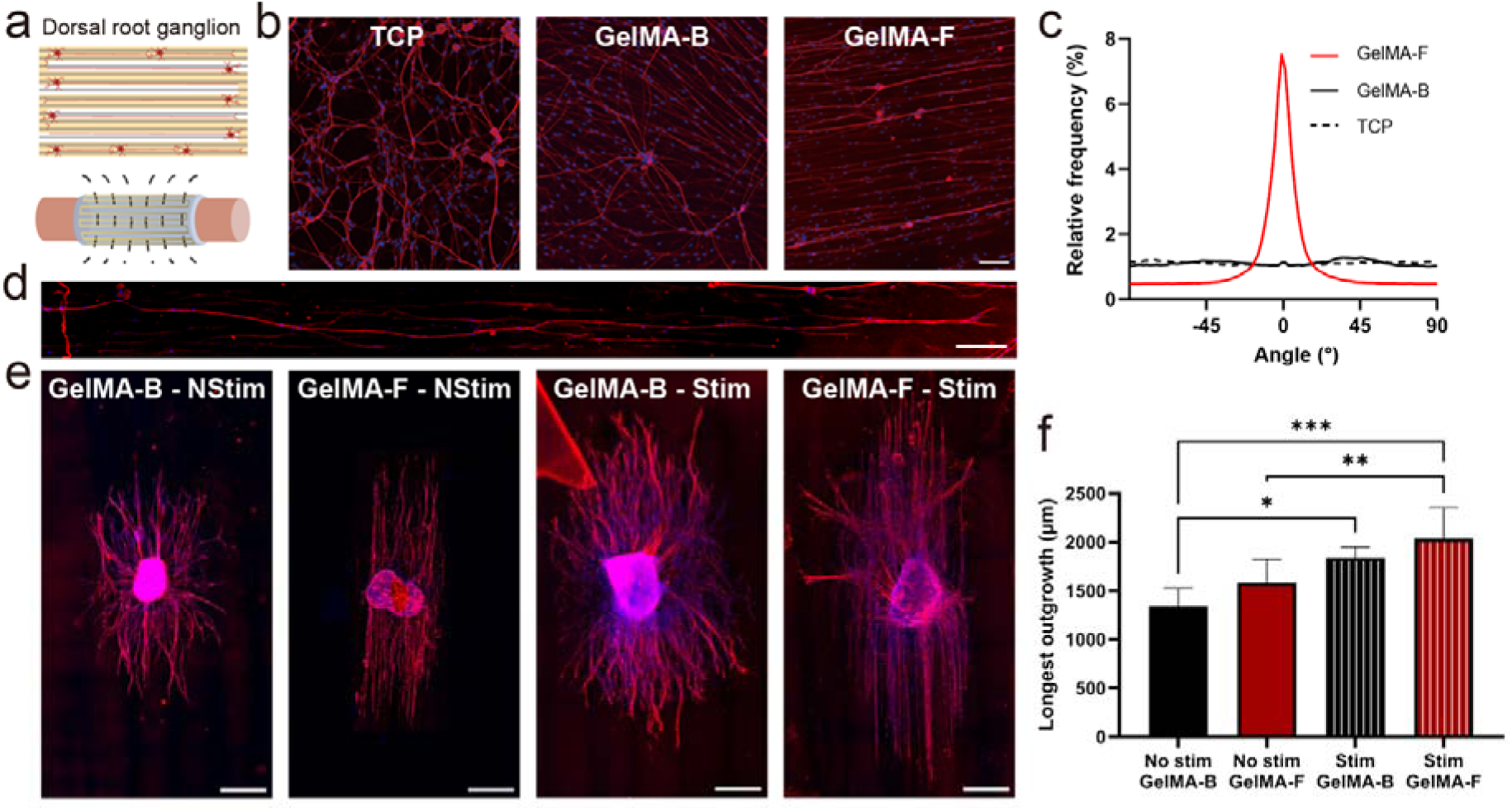
GelMA-F guided and EM stimulated outgrowth of DRG neurons. **a)** Illustration of GelMA-F as a nerve guide conduit with wireless EM stimulation**. b)** Dissociated DRG neurons after 7 days of culture on TCP, on the surface of GelMA-B and GelMA-F. Red = β-III-tubulin, blue = Hoechst. Scale bar 100 µm. **c)** Directionality analysis of DRG axons on TCP, GelMA-B, and GelMA-F after 7 days of culture. **d)** A stitched image of DRG neuron axons after 7 days of culture on GelMA-F. Scale bar 200 µm. **e)** Fluorescence images of DRG explants on 4% w/v GelMA-F after 7 days of culture with wireless EM stimulation. Red = β-III-tubulin, blue = Hoechst. Scale bar 500 µm. **f)** Longest outgrowth quantification of DRG explants after 7 days of culture.

DRG explants were cultured on EM patches encapsulated in GelMA-B or GelMA-F for 7 days with EM stimulation (7.2 mV, 90 kHz, 2 h/day, 7 days) (**Figure 6.e**). Evident aligned neural outgrowth along the filamented GelMA-F surface topography is observed. (**Figure S5**) Sholl analysis for longest outgrowth revealed a significant increase in outgrowth length in EM stimulated samples compared to non-EM stimulated samples (**Figure 6.f**).

The successful fabrication of GelMA-F EM nerve guide conduit and subsequent neurite outgrowth along the fibers indicate the promise of its use for peripheral neuron repair. The ease at which tubular structures of oriented hydrogel fibers can be bioprinted within seconds makes it a promising fabrication technique for solving challenges such as nerve misalignment, which leads to incomplete reinnervation and slows the recovery of muscle function post nerve repair [74, 75]. Potentially, a personalized nerve guide conduit may be rapidly produced within seconds to match the specific nerve bundle structure, increasing the likelihood of joining together the correct nerve fibers in a damaged nerve bundle.

### Stimulated Anisotropic Hydrogels Drive Comprehensive and Functional Nerve Regeneration *In Vivo*

To challenge the existing limits of peripheral nerve repair, we deployed our integrated system in a critical-sized 1 cm rat sciatic nerve defect. To prepare GelMA-F nerve conduits, GelMA-F together with EM coil was carefully inserted into an electrospun PCL nanofibrous tube (PCL). All conduits demonstrated excellent *in vivo* biocompatibility over the 8-week period, with no signs of severe adverse events or systemic toxicity (**Figure S6**). Our findings reveal that the seamless integration of anisotropic guidance and wireless EM stimulation orchestrated a regenerative cascade of remarkable fidelity. This process initiated the de novo construction of a native-like nerve architecture, was sustained by a meticulously sculpted pro-regenerative microenvironment, and culminated in a degree of functional recovery that rivals the clinical gold standard. We will now dissect this comprehensive success by following the evidence presented in **Figure 7**.

**Figure 7.**
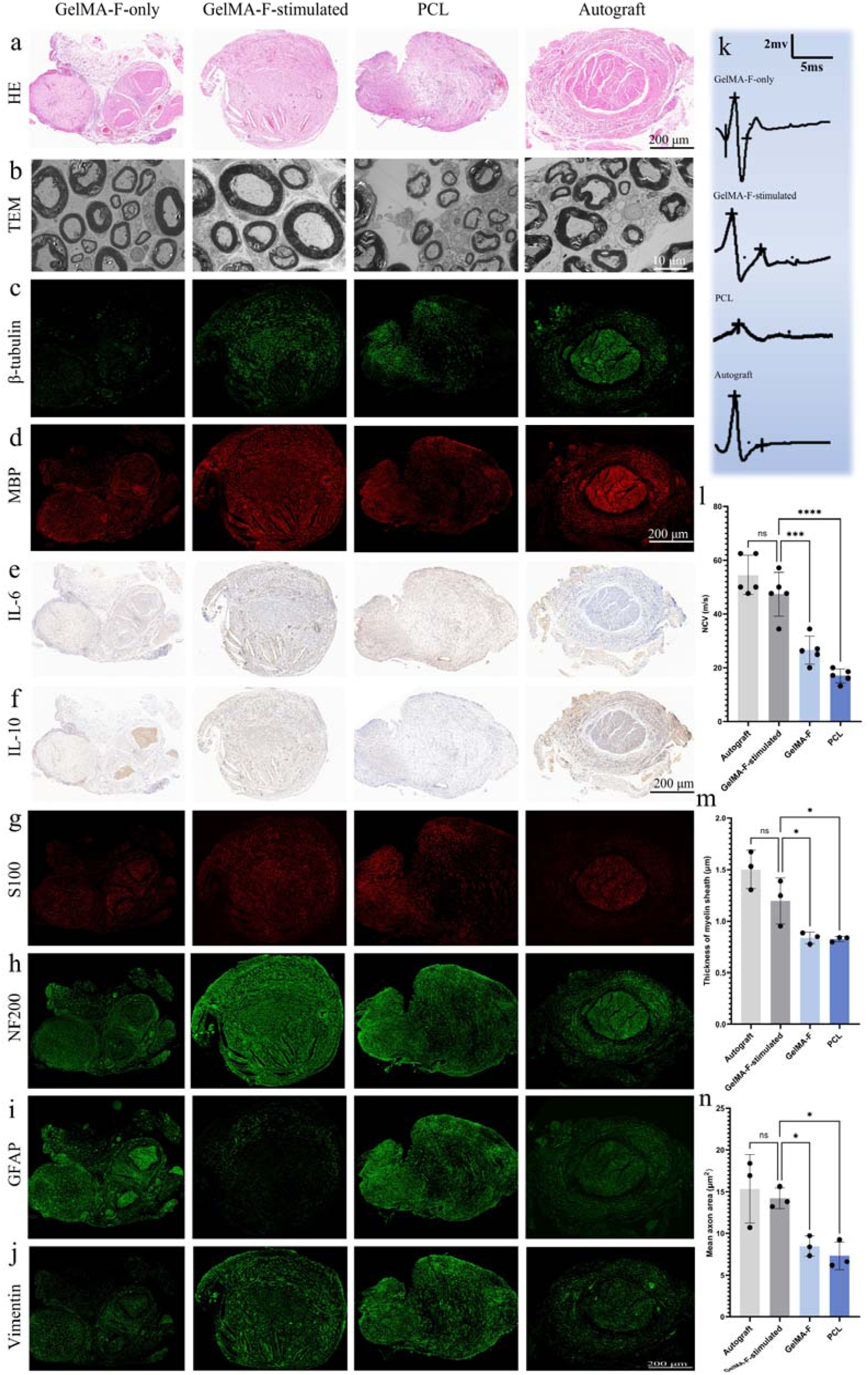
Anisotropic Hydrogels with Wireless Stimulation Drive Comprehensive Nerve Regeneration and Functional Recovery In Vivo. All data were collected 8 weeks post-implantation in a 1 cm rat sciatic nerve defect model. The GelMA-F-stimulated group is compared against GelMA-F (non-stimulated), PCL-only conduit, and Autograft controls. (**a-d, m, n**) Structural and ultrastructural restoration of regenerated nerve. (**a**) HE staining demonstrating the density and organization of regenerated nerve fibers. (**b**) TEM revealing thick, organized myelin sheaths, a hallmark of high-quality remyelination. (**c, d**) Immunofluorescence for (**c**) the axonal marker β-tubulin (green) and (**d**) the myelin marker MBP (red), showing robust axonal growth and myelination. (**m, n**) Quantification confirms that (**m**) myelin sheath thickness and (**n**) mean axon area in the stimulated group achieved metrics statistically comparable to the autograft. (**e-j**) Engineering a pro-regenerative microenvironment. (**e, f**) Immunohistochemistry shows (**e**) reduced pro-inflammatory IL-6 and (**f**) elevated anti-inflammatory IL-10. (**g-j**) Immunofluorescence reveals (**g**) abundant S100-positive Schwann cells, (**h**) mature NF200-positive axons, (**i, j**) suppressed GFAP and Vimentin expression, indicating reduced fibrosis. (**k, l**) Restoration of physiological function. (**k**) Representative evoked nerve action potentials and (**l**) quantification of nerve conduction velocity (NCV) demonstrate that the GelMA-F-stimulated group restored signal transmission to near-autograft levels. Scale bars: 200 µm (a, c, d, g-j); 10 µm (b). Data in (**k-m**) are presented as mean ± SEM (n =3). Statistical significance was assessed by one-way ANOVA with Tukey’s post-hoc test. ns, not significant; *P < 0.05, **P < 0.01, ***P < 0.001.

The cornerstone of this regeneration was the system’s ability to guide the growth of nerve tissue with remarkable structural integrity. Histological analysis revealed that the GelMA-F-stimulated group fostered densely packed, highly organized axons within a well-defined structure, closely resembling the architecture of the autograft gold standard (**Figure 7a**, HE). This observation was confirmed at the ultrastructural level, where TEM images displayed axons encased in thick, uniform myelin sheaths—a hallmark of successful and high-quality remyelination (**Figure 7b**). Immunofluorescence staining further substantiated these findings, showing the most extensive axonal formation via strong β-tubulin expression (**Figure 7c**) and the highest degree of myelination through intense myelin basic protein (MBP) staining (**Figure 7d**) in the GelMA-F-stimulated group.

This robust structural repair was supported by a meticulously engineered, pro-regenerative microenvironment [76]. Adequate blood supply is critical for nerve repair [77], while CD31 and CD34 staining demonstrated that our system effectively promoted angiogenesis around the lesioned nerve (**Figure S7**). Moreover, the system actively sculpted a favorable immune milieu by suppressing the pro-inflammatory cytokine IL-6 (**Figure 7e**) while elevating the anti-inflammatory cytokine IL-10 (**Figure 7f**). This anti-inflammatory state was crucial for subsequent cellular processes [78]. Indeed, it supported a thriving population of S100-positive Schwann cells (**Figure 7g**), which are essential for myelination and trophic support. This, in turn, promoted the development of mature, large-diameter axons, as indicated by strong NF200 signals (**Figure 7h**). Critically, the system potently inhibited scar formation, evidenced by markedly reduced expression of the fibrotic and reactive glial marker GFAP and Vimentin, both of which also represents the activated Schwann cells (**Figure 7i-j**). This anti-fibrotic action created a permissive, barrier-free pathway essential for axonal elongation [79].This robust structural repair was supported by a meticulously engineered, pro-regenerative microenvironment [76]. Adequate blood supply is critical for nerve repair [77], while CD31 and CD34 staining demonstrated that our system effectively promoted angiogenesis around the lesioned nerve (**Figure S7**). Moreover, the system actively sculpted a favorable immune milieu by suppressing the pro-inflammatory cytokine IL-6 (**Figure 7e**) while elevating the anti-inflammatory cytokine IL-10 (**Figure 7f**). This anti-inflammatory state was crucial for subsequent cellular processes [78]. Indeed, it supported a thriving population of S100-positive Schwann cells (**Figure 7g**), which are essential for myelination and trophic support. This, in turn, promoted the development of mature, large-diameter axons, as indicated by strong NF200 signals (**Figure 7h**). Critically, the system potently inhibited scar formation, evidenced by markedly reduced expression of the fibrotic and reactive glial marker GFAP (**Figure 7i**). This anti-fibrotic action created a permissive, barrier-free pathway essential for axonal elongation [79].

Ultimately, this comprehensive structural and microenvironmental restoration translated into definitive functional recovery. Electrophysiological measurements revealed robust evoked neural signals in the GelMA-F-stimulated group, with amplitudes comparable to the autograft (**Figure 7k**). Consequently, the nerve conduction velocity (NCV) reached 47.36 m/s in the GelMA-F-stimulated group, significantly outperforming controls and closely approaching the gold standard (**Figure 7l**). This functional restoration was directly corroborated by the superior quantitative metrics of the regenerated structure, including a myelin sheath thickness (**Figure 7m**) and mean axon area (**Figure 7n**) that were statistically equivalent to the autograft. Furthermore, this successful reinnervation effectively prevented atrophy of the target gastrocnemius muscle (**Figure S8**), providing the ultimate proof of a restored nerve-to-muscle connection.

In summary, the therapeutic success detailed above is not merely an additive effect but a direct result of the powerful synergy between physical guidance and electrical stimulation. While the anisotropic conduit provided the essential “physical highway” for organized growth, the wireless stimulation acted as a “biological accelerator,” modulating the immune environment, promoting crucial cell populations, and enhancing the overall pace and quality of regeneration. It is this integrated, synergistic strategy that enabled the system to overcome the complex barriers of nerve repair and achieve a therapeutic outcome comparable to the clinical gold standard.

### Conclusions

In conclusion, our findings reveal that the soft, anisotropic architecture of biohybrid system exerts powerful topographical control over neural morphogenesis across central and peripheral systems. By selectively promoting neuronal rather than glioblastoma-derived outgrowth, these materials establish a microenvironment conducive to regeneration rather than malignancy, offering a new strategy for re-establishing functional neural circuits in the central nervous system. When combined with wireless EM stimulation, the system provides dynamic electric modulation that further enhance hippocampal neural network integration and peripheral neurite extension. The resulting synergy between structural guidance and remote stimulation culminates in *in vivo* regeneration outcomes that approach the efficacy of clinical autografts. This work establishes a multiscale biohybrid strategy in which the synergy between soft biomaterials and programmable wireless bioelectric cues enables noninvasive regulation of neural morphogenesis, defining key design principles for future bioelectronic interfaces and regenerative scaffolds.

## Methods and materials

### Materials and fabrication of the EM stimulation patch

The fabrication of the EM receiver coil follows the procedure described in our previous paper with minor alterations [16]. Polycaprolactone (PCL) was used as raw material and was electrowritten without any modifications (medical grade, PURESORB PC 08 PCL, Corbion). The PCL receiver coil on PCL support grids were electrowritten using a melt electrowriting instrument (CAT000111, Spraybase, Ireland). The structure was designed and written in G-code which was run by a Mach3 motion control software. PCL was melted at 80 °C for one hour before printing through a stainless-steel needle (24 G). A 10 cm diameter Silicon-wafer was used as a collector at a 4 mm distance from the needle. A 4.25-4.5 kV high voltage was applied between the needle and the collecting plate. A flow rate of 0.55 bar was controlled by a Spraybase® Pressure Driven System. The coil (EM receiver) was printed at a rate of 1000-1050 mm min^−1^ with a support grid at a rate of 75 mm min^−1^. The coil structure and support grid were printed with 40 layers and 2 layers, respectively. The G-code used for melt electrowriting of the anisotropic neural stimulation patch can be found in Supplementary Table 1. The resulting patch was subsequently coated with 4 nm titanium and 80 nm of gold using E-beam GLAD (Cryofox Explorer 500 GLAD). During coating the scaffolds were tilted 20 degrees and rotated at 3 rpm.

The wireless EM stimulation system used for stimulating neural cell growth is comprised of a receiver coil patch placed in a 24-well plate and a rectangular transmitter coil placed around the well plate. The transmitter coil size is matched to fit around a standard well plate to minimize the distance between the patch and the transmitter coil. The transmitter coil was made of 100 turns of copper wire (11 cm x 15 cm) connected to a Waveform Generator (HP 33120A, Hewlett Packard) supplying alternating current. The voltage and frequency applied through the transmitter coil were adjusted to produce the desired voltage output in the receiver coil scaffold.

### GelMA synthesis

Gelatin type A from porcine skin was dissolved at 10% w/v in PBS buffer pH 7.2 (Gibco) at 60 °C under stirring overnight. Methacrylic anhydride (MA) was added dropwise at a rate of 0.5 mL/min to a final MA concentration of 10% v/v using a syringe pump under vigorous stirring. The GelMA solution was allowed to react for 3 hours at 50 °C under stirring. The GelMA solution was diluted with 50 °C PBS pH 7.2 to one-fifth the concentration under stirring and subsequently transferred to dialysis tubes and dialyzed against 10x volume distilled water at 50 °C for 7 days with water changes every day. The dialyzed GelMA solution was freeze dried for 7 days and stored at −20 °C until use.

### Material characterization

The voltage output of the receiver coil was recorded by an Agilent DSOX3012A oscilloscope.

The compression stress-strain measurements of GelMA were performed by compression of cylinders (Ø=8.4 mm, L=20 mm) of freshly prepared 2%, 3%, and 4% w/v GelMA using a BOSE Electroforce 3200 (BOSE). The cylinders were compressed at a rate of 1 mm/s with a force resolution of 0.1g.

### Preparation of projection images for bioprinting

The projection images were created with Inkscape, and their dimensions were set to a fixed resolution of 1024 x 768 pixels. The images were in 8-bit greyscale, where completely white pixels represented full light projection. The width of each pixel was set by the system to approximately 27 µm. The files can be found in supplementary information **(Figure S9)**.

### GelMA-B and GelMA-F fabrication

To prepare GelMA precursor, freeze dried GelMA was dissolved in PBS at 37 °C for 1 hour to the desired concentration. A stock solution of lithium phenyl-2,4,6-trimethylbenzoylphosphinate (LAP, Sigma-Aldrich) (50 mg ml^-1^) was added to achieve a final concentration of 1 mg ml^-1^ LAP. The GelMA precursor was filtered by a 0.2 µm filter before use to sterilize and remove particles. For encapsulation of eGFP-MSC cells in GelMA-B and GelMA-F, eGFP-MSC cells were resuspended in GelMA precursor before light exposure at a cell density of 0.4 x 10^6^ cells ml^-1^.

To fabricate GelMA-F structures, the GelMA precursor was transferred to sterile glass vials (Ø = 1.5 cm) and was thermally crosslinked in a 4 °C water bath for 30 minutes. The image projections were performed using a tomographic volumetric printer (Tomolite version 2.0 Performance-closed, Readily3D SA). The light intensity was set to 10.5 mW cm^-2^. For EM coil integrated GelMA-F, the EM coil was inserted in the printing vessel with its long walls align with the beam projection. After projection, the photo-crosslinked GelMA was reheated to 37 °C in a water bath and the structure was retrieved with a sterile spatula. Non-crosslinked GelMA washed away with PBS and replaced with warm growth media for cell laden structures.

To fabricate GelMA-B, the GelMA precursor was transferred to a Ø=5cm petridish and crosslinked utilizing a 365 nm UV lamp (6W, VL-6.L, Vilber Lourmat). The structure was then cut, transferred to culture wells using a sterile spatula, rinsed with 37 °C and replaced with warm growth media for cell laden structures.

### eGFP-MSC Cell culture

eGFP-MSC cells were precultured in a T75 culture flask (Sarstedt) in growth medium (10% fetal bovine serum (FBS), 1% penicillin-streptomycin (P/S) in high glucose DMEM (Gibco)). Prior to GelMA-B and GelMA-F biofabrication, eGFP-MSC cells in a T75 culture flask at ∼80% confluency was rinsed twice with prewarmed PBS (37 °C), and trypsinized with 0.05% Trypsin-EDTA (Gibco) for 5 minutes at 37 °C. The trypsinization was inhibited with twice the volume of growth medium and centrifuged for 5 minutes at 250g. The cells were resuspended in fresh growth medium or GelMA precursor and further used as described in the GelMA-B and GelMA-F biofabrication method section. Growth media was fully changed every 2-3 days.

### DRG dissection and culturing

Neuronal-enriched DRG cultures were prepared as previously described with the following modifications [80]. Postnatal day 1-4 C57BL/6JRj mice were euthanized and disinfected in 70 % ethanol. 40 DRGs from each mouse were dissected and pooled into ice-cold HBSS medium (STEMCELL Technologies, 37150). The DRGs were used as whole DRGs (DRG explants) or the DRGs were dissociated and cultured as neuron enriched DRG cultures (DRG neurons).

For experiments using DRG explants, the DRGs were cut in half to open the protective DRG membrane and stored in ice cold DMEM (Gibco, 41965-039) with 10 % (v/v) FBS (Sigma-Aldrich, F9665) and 1 % (v/v) pen/strep (Sigma-Aldrich, P4333) until seeded onto pre-prepared 4% GelMA-F or GelMA-B structures. The DRG explants were maintained in a humidified atmosphere at 37°C and 5% CO_2_ in DRG culture medium (DMEM with 10 % (v/v) FBS, 1 % (v/v) pen/strep, 25 ng/ml neurotrophin-3 (NT-3, Alomone Labs, N260), and 25 ng/ml nerve growth factor (NGF, 2.5S, Gibco, 13257-019)) for 7 days. Half the volume of growth medium was replaced with fresh DRG culture medium every 2 days.

For dissociated DRG cultures, the DRGs from 4 pups were centrifuged at 500xg for 4 min at room temperature. DRGs were then enzymatically dissociated for 30 min at 37°C, 5% CO_2_ using 13.33 U/ml papain (Worthington, LS003126). The DRGs were then centrifuged for 1 min at 500xg, papain was aspirated, and they were further digested for 30 min in 2.5 mg/ml collagenase (Worthington, LS004176) and 5 U/ml dispase II (Sigma-Aldrich, D4693) at 37°C, 5% CO_2_. The digested tissue was centrifuged for 1 min at 500xg and the collagenase/dispase solution was aspirated. The tissue was washed in DMEM with 10 % (v/v) FBS and 1 % (v/v) pen/strep, centrifuged for 1 min at 500xg and mechanically dissociated by gently trituration using a p1000 pipette in the same buffer with 40 Kunitz units Deoxyribonuclease I (Sigma-Aldrich, DN25-1G). Debris was removed from the cell suspension by filtering through a 100 µm cell strainer (Falcon, 352360). Dissociated cells were pre-plated on non-coated plates for 1.5 hours at 37°C, 5% CO_2_. The neuron-enriched media was subsequently collected and centrifuged for 1 min at 500xg. The cell pellet was resuspended in DRG culture media (DMEM with 10 % (v/v) FBS, 1 % (v/v) pen/strep, 25 ng/ml neurotrophin-3, and 25 ng/ml nerve growth factor. The cell suspension was then seeded onto 4% w/v GelMA-B or GelMA-F, or onto poly-D-lysine+laminin coated TCP at a seeding density of 50 × 10^3^ cells/cm^2^. The DRG cells were maintained in a humidified atmosphere at 37°C and 5% CO_2_ in DRG culture medium for 7 days. Half the volume of growth medium was replaced with fresh DRG culture medium every 2 days.

### Hippocampal neuron cell culture

Dissociated rat hippocampal neuron cultures were prepared and maintained as previously described [81]. Briefly, rat hippocampi were dissected from postnatal day 0 to 1 pups of either sex (Sprague-Dawley strain; Charles River Laboratories), dissociated with papain (Sigma-Aldrich), and then seeded at a density of 40 × 10^3^ cells/cm^2^ onto 4% w/v GelMA-F or poly-d-lysine-coated tissue culture plates (SARSTEDT). Neurons were maintained in a humidified atmosphere at 37°C and 5% CO_2_ in hippocampal growth medium (Neurobasal-A supplemented with B27 and GlutaMAX-I; Life Technologies) for 21 days *in vitro*. Half the volume of hippocampal growth medium was replaced with fresh hippocampal growth medium twice per week.

### Glioblastoma tissue acquisition

The glioblastoma tissue acquisition followed the procedure described in our previous work [82]. All procedures with human tissue and data were approved by and conducted in accordance with the Scientific Ethics Committee for the Region of Midtjylland Denmark (official name in Danish: De Videnskabsetiske Komitéer for Region Midtjylland) (journal number: 1-10-72-82-17) and conducted in accordance with the ethical principles of the World Medical Association Declaration of Helsinki [83]. Surgical specimens were collected at the neurosurgical department at Aarhus University Hospital (Denmark) through collaboration with local surgeons. Before participating in the study, all patients gave informed consent for the donation of tissue. The patients included in the study were undergoing surgery for GBM, and the obtained specimens were surgically excised cancer-infiltrated cortical tissue that required removal as part of the surgical treatment. The patients’ diagnoses were confirmed based on pathology examination and standardized WHO criteria [84].

Preoperatively, the patients were administered the fluorescent compounds 5-Aminolevulinic acid hydrochloride (5-ALA) (20 mg/kg bodyweight administered orally) and sodium fluorescein (200 mg IV), which allowed the neurosurgeon to visualize the tumor and the areas infiltrated by the tumor [85, 86]. The surgical specimens, ranging between 1-2 cm^3^ in volume, were resected from the superficial cortical regions and passed the contrast-enhancing rim of the tumor. The procedure did not impose any constraints on the patients or their surgical treatment. The excised specimens were transferred immediately after resection to a sterile container filled with slicing artificial cerebrospinal fluid (ACSF). The slicing ACSF comprised (in mM): 75 Sucrose, 84 NaCl, 2.5 KCl, 1 NaH_2_PO_4_, 25 NaHCO_3_, 25 D-glucose, 0.5 CaCl_2_·2H_2_O and 4 MgCl_2_·6H_2_O. The pH was adjusted to 7.3–7.4 using concentrated hydrochloric acid. The osmolality was confirmed to be between 295– 305 mOsm kg^−1^. The solution was pre-chilled and thoroughly bubbled with carbogen (95% O2, 5% CO2) gas before tissue collection. During transportation, the tissue was kept in a continuously carbogenated slicing ACSF solution maintained at a constant temperature. The duration of tissue transport from the operating room to the laboratory site ranged from 15 to 30 minutes.

### Tissue processing

The tissue processing followed the procedure described in our previous work [82, 87]. The human ex vivo brain slice culture method was developed based on previously established procedures [88–90]. The tissue samples underwent thorough examination to identify the most effective method for blocking and mounting them onto the vibrating microtome platform, Leica 1200S vibratome (Leica Microsystems, Denmark). The samples were sliced perpendicular to the pial surface to preserve dendrite and axon integrity in the GBM-infiltrated region. The tissue slicing procedure was performed in sterile-filtered and chilled ACSF slicing solution. The solution was saturated with carbogen gas to maintain tissue oxygenation throughout the processing steps. Slices of 300 μm thickness were cut and transferred to a carbogenated (95% O2/5% CO2) and tempered (34 °C) slicing ACSF solution to recover for 30 min. The slices were then transferred onto membrane inserts (Millipore) in 6 well plates with 800µL per well of slice culture media of the composition: 8.4 g/L MEM Eagle medium, 20% heat-inactivated horse serum, 30 mM HEPES, 13 mM D-glucose, 15 mM NaHCO_3_, 1 mM ascorbic acid, 2 mM MgSO_4_·7H_2_O, 1 mM CaCl_2_.4H_2_O, 0.5 mM GlutaMAX-I and 1 mg/L insulin, 25 U/mL Penicillin/Streptomycin. The slice culture medium was adjusted to pH 7.2–7.3 by addition of 1 M Tris base. The osmolality was adjusted to 295–305 mOsmoles kg^−1^ by addition of pure H_2_O. The tissues were cultured humidified with 5% CO_2_ in an incubator at 37°C. The slice culture medium was replenished every one to two days until the experiment was completed. One to three hours after the brain slices were transferred onto cell culture inserts, the brain slices were transduced by direct application of concentrated AAV viral particles over the slice surface for fluorescent labelling. Two days later the slices were transferred onto 1 mm thick slices of 2% w/v GelMA-B or GelMA-F on membrane inserts, or directly onto separate membrane inserts in 6-well plates.

### AAV-viral transduction

The adeno-associated virus (AAV) transduction followed the procedure described in our previous work [82, 87]. The AVV vectors were acquired from the University of Zurich Viral Vector Facility. The optimized hGFAP-AAV vector virus (ssAAV5-hGFAP-hHEbl/E-EGFP-bGHp(A)) was used to label astrocyte or glioblastoma cells with fluorescent eGFP. The functional virus titers ranged between 1–3 × 10^13^ units per mL. Human slice cultures were transduced with AAV amplicons by directly applying the concentrated viral stock solutions to the slice surface using a fine micropipette. This was done at least one hour after plating to allow the slice to equilibrate. The virus was carefully spread over the entire slice surface, ensuring that the surface tension of the viral solution was maintained without it spreading onto the membrane insert.

### Live/dead imaging

eGFP-MSC cells were cultured and maintained as described in the Cell Culture section. After the stated days of culture, the sample wells were gently rinsed with warm growth media. Live/dead solution was prepared by diluting Calcein-AM (2 mM in DMSO) 1:1000 and propidium iodide (PI) (1.5 mM in deionized H_2_O) 1:500 in 37 °C cell culture medium without serum and phenol red. The growth medium was removed. 300 µL Live/dead solution was added to each sample well and incubated for 30 minutes at 37 °C and 5% CO_2_ protected from light. The live/dead solution was subsequently removed and replaced with 37 °C cell culture medium without serum or phenol red. Fluorescence imaging was immediately performed using EVOS FL Auto Cell Imaging System (Invitrogen). For Calcein 494nm/517nm (excitation/emission) was used and PI 535nm/617nm was used.

### Calcium imaging

Hippocampal neurons were cultured and maintained as described in the cell culture section. After the stated days of culture the samples were rinsed with warm modified Hanks Balanced Salt Solution (HBSS) (3.26 mM CaCl_2_, 0.492 mM MgCl_2_, 0.406 mM MgSO_4_, 5.33 mM KCl, 0.441 mM KH_2_PO_4_, 4.17 mM NaHCO_3_, 138 mM NaCl, 0.336 mM Na_2_HPO_4_, 20 mM D-glucose). The samples were incubated with 1 µM Oregon Green 488 BAPTA-1 (OBG-1 AM, Invitrogen, O6807) and 0.025% Pluronic F-127 (Sigma-Aldrich, P2443) for 60 minutes at 37 °C and 5% CO_2_ protected from light. The calcium staining solution was subsequently removed, rinsed with 37 °C HBSS, and further incubated for 60 minutes at 37 °C and 5% CO_2_ protected from light in HBSS. Fluorescence calcium imaging using 494nm/517nm (excitation/emission) at 37 °C and 5% CO_2_ was performed using EVOS FL Auto Cell Imaging System (Invitrogen) with a built-in incubator. Videos were taken at 20x magnification and 20 fps and stored as individual 16-bit greyscale TIF format images. Image and data analysis was performed using ImageJ with the Spikey plugin.

### Immunostaining and fluorescence imaging

After ended culture, samples were fixed in 4% formaldehyde solution for 20 minutes, and subsequently permeabilized in PBS with 2% Triton X-100 for 15 minutes. After blocking in 1% bovine serum albumin (BSA)/PBS (Sigma, A2153) solution for 1 hour at room temperature cells were immunostained with rabbit anti-β-III-tubulin (Abcam ab18207, 1:1000) primary antibody, chicken anti-GFAP (Invitrogen, PA1-10004, 1:1000), and Guinea pig anti-Synaptophysin (Alomone Labs, ANR-013-GP200UL, 1:200) at 4 °C overnight. After washing with PBS, cells were stained with Donkey anti-rabbit (Alexa Fluor 594, Abcam ab 150076, 1:1000), Goat anti-chicken IgY (Alexa Fluor TM 488, Invitrogen, A-11039, 1:500), and goat anti-guinea pig IgG (Alexa Fluor 633, Invitrogen, A-21105, 1:1000) secondary antibody for 1 hour at room temperature. After washing with PBS again, Hoechst 33258 (Life Technologies, 1:10,000) was added for 10 minutes to stain cell nuclei. For brain tissue slices the permeabilization, blocking, staining, and washing steps were performed using Perm/Wash (BD Biosciences, 554723) instead of Triton X-100, BSA, BSA, and PBS, respectively. Fluorescence images were acquired with an EVOS FL Auto Cell Imaging System (Invitrogen).

Cell neurite length was manually traced and evaluated from cell fluorescence images using ImageJ [91] with NeuronJ [92] plugin-software. Neurites longer than the cell body and not extending outside of the field of view were included. The average neurite length is defined as the total neurite length divided by the total neurite number. At least 300 neurites were measured from each condition. The manual tracing was performed blinded to the experimental condition.

Sholl analysis was performed by first thresholding images of β-III-tubulin stained DRGs and subsequently performing automated Sholl analysis by using the neuroanatomy-Sholl plugin in ImageJ. The longest neurite was extracted from each sample as the farthest point from the center of Sholl circle. At least 6 samples were measured for each condition.

### Nerve Conduit Fabrication

PCL (PansiTech) was dissolved in 1,1,1,3,3,3-Hexafluoro-2-propanol (HFIP) at a concentration of 12% (w/v). The mixture was continuously stirred at 60 °C and a rate of 500 rpm for 1 hour, followed by 1 hour of ultrasonication to ensure complete dissolution and dispersion. The resulting PCL solution was electrospun into fibrous membranes, and then were cut and subsequently rolled into tubular conduits, each measuring approximately 1 cm in length with an outer diameter of 0.6 cm. To prepare GelMA-F nerve conduits, pre-fabricated GelMA-F hydrogel scaffolds were carefully inserted into the PCL tubes. Both GelMA-F nerve conduits and PCL-only conduits were sterilized by UV irradiation in a sterile laminar flow hood before in vivo implantation.

### Animal Model and Surgical Implantation

All animal procedures were approved by the Institutional Animal Care and Use Committee of the Shanghai Jiaotong University affiliated Shanghai Sixth People’s Hospital. A total of 20 male Sprague-Dawley (SD) rats (12 weeks old; 210–250 g) were used and randomly allocated to 3 groups: autografted group (n = 5), GelMA-F group (n = 10) and PCL group (n = 5). Briefly, the rats were first anesthetized with sodium pentobarbital and a skin incision was made in the posterolateral aspect of the right mid-thigh. The biceps femoris muscle was carefully separated to fully expose the sciatic nerve which was then completely transected, creating a 1 cm segmental defect. The GelMA-F nerve conduit was interposed into the 1 cm defect, and a 6-0 angle needle was applied to carefully suture both the proximal and distal nerve stumps to the conduit ends. Subsequently, the biceps femoris muscle and skin were closed in turn using 4-0 absorbable sutures. After operation, each rat received a penicillin injection to prevent postoperative infection. Similar surgical procedures were performed for rats receiving PCL conduits and autologous nerve grafts, serving as control groups.

### In Vivo EM Stimulation and Tissue Processing

Rats implanted with GelMA-F nerve conduits were randomly divided into two subgroups (n=5 per subgroup): a stimulated group and a non-stimulated group. The stimulated subgroup received wireless EM stimulation (4 mT intensity, 30 min/session) every other day. Eight weeks post-surgery, all animals were sacrificed and the regenerated sciatic nerve segments at the defect site were carefully harvested. Additionally, the gastrocnemius muscles from both ipsilateral and contralateral hind limbs, along with major organs, were collected for subsequent analysis.

### Electrophysiological Assessment

Prior to tissue harvest, sciatic nerve functional recovery was assessed through electrophysiological measurements. Rats were deeply anesthetized with an intraperitoneal injection of 50 mg/kg pentobarbital sodium solution. The skin and biceps femoris muscle over the sciatic nerve were carefully dissected to expose the regenerated nerve segment. Nerve conduction velocity (NCV) measurements were then performed on the exposed regenerated nerve to quantify functional recovery.

### Masson Staining

Gastrocnemius muscles were immediately put into 4% paraformaldehyde (Servicebio, G1101) for 24h. Muscles were taken out of 4% paraformaldehyde and smoothed using the scalpel in the fume hood. The cut muscles and corresponding labels were put in the embedding frame, and then, the dehydration box were put into the dehydrator (DIAPATH, Donatello) in order to be dehydrated with gradient alcohol: 75% alcohol for 4 h,85% alcohol for 2 h,90% alcohol for 2 h,95% alcohol for 1 h, anhydrous ethanol I for 30 min, anhydrous ethanol II for 30 min, alcohol benzene for 10 min, xylene II for 10 min, 65℃ melting paraffin I for 1h, 65℃ melting paraffin II for 1h, 65℃ melting paraffin III for 1 hour. The wax-soaked muscles were embedded in the embedding machine (Junjie Eletronics Co., Ltd, JB-P5). The trimmed wax blocks were put into a paraffin slicer (Leika Instrument Co., Ltd, RM2016) for slicing, with a thickness of 4 μm. Slices were flattened when they floated on the 40 ℃ warm water of the spreading machine (Kehua Instrument Co., Ltd, KD-P), and slices were picked up by the glass slides and baked in the oven at 60 ℃. After the water-baked dried wax melted, it was taken out and stored at room temperature.

The paraffin sections were immersed in sequence in environmentally friendly dewaxing transparent liquid I (Servicebio, G1128) for 20 min - environmentally friendly dewaxing transparent liquid II for 20 min - anhydrous ethanol I for 5 min - anhydrous ethanol II for 5 min - 75% ethyl alcohol for 5 min, and then rinsed with tap water. The frozen muscle slices were removed from the −20℃ freezer and restored to room temperature, fixed with 4% paraformaldehyde (Servicebio, G1101) for 15 min, and then rinsed with running water. The slices were soaked in masson A (Servicebio, G1006) overnight, rinsed with tap water. masson B and masson C were prepared into masson solution according to the ratio of 1:1, and then, stained with masson solution for 1 min, rinsed with tap water, and differentiated with 1% hydrochloric acid alcohol (Servicebio, G1039) for several seconds, rinsed with tap water. The slices were soaked in masson D for 6 min, rinsed with tap water, and, soaked in masson E for 1 min. Slices were slightly drained and soaked in masson F for 30 s. The slices were rinsed with 1% glacial acetic acid and then dehydrated with two cups of anhydrous ethanol. Slides were soaked in 100% ethanol for 5 min, xylene for 5 min, and finally sealed with neutral gum. Slides were inspected with microscope, and images were acquired and analyzed.

### TEM

Nerve samples were immediately fixed in 2.5% glutaraldehyde (Servicebio G1102) after excision, and then processed (embedded and sectioned to approximately 0.5 μm thickness) following the masson staining protocol. Ultrathin sections were then double-stained with uranyl acetate and lead citrate, and subsequently examined and photographed with TEM (HITACHI, HT7700).

### Histological, Immunohistochemical, and Immunofluorescence Analyses

The heart, liver, spleen, lung, kidney and nerves were collected and immediately fixed in 4% paraformaldehyde (Servicebio, G1101) for 24 h. Paraffin sections were prepared and frozen sections were fixed following the masson staining protocol. Slices were put into hematoxylin solution (Servicebio G1003) for 3 min, rinsed with tap water, and then, differentiated with hematoxylin differentiation solution and rinsed with tap water. Then slices were treated with hematoxylin bluing solution, rinsed with tap water, and then placed in sequence in 85% ethanol for 5 min - 95% ethanol for 5 min-eosin dye for 5 min. Slices were put into absolute ethanol I for 5 min-absolute ethanol II for 5 min-absolute ethanol III for 5 min-xyleneⅠfor 5 min-xyleneⅡfor 5 min, and then, were sealed with neutral gum. Slices were stained by immersion in toluidine blue (Servicebio, G1032) for 5 min, and then, were briefly differentiated using 0.1% glacial acetic acid (SCRC,1000218) and terminated by rinsing with tap water. After air-drying, slices were immersed in xylene (SCRC,10023418) for 10 min and sealed with neutral gum. Slides were inspected with microscope to acquire and analyze HE staining images.

Before immunohistochemical and immunofluorescence staining, antigen retrieval was performed on neural slices using microwave heating in citrate buffer (pH 6.0). After blocking with 3% BSA (Servicebio, GC305010) in PBS (Servicebio, G0002) for 30 min at room temperature, slices were incubated overnight at 4 °C with rabbit anti-β-III-tubulin (Servicebio, gb15139, 1:600), rabbit anti-MBP (Servicebio, gb11226-1, 1:500), rabbit anti-S100 (Servicebio, anti-CD31 (Servicebio, GB11063-2, 1:600), rabbit anti-CD34 (Servicebio, GB15013, 1:500), rabbit anti-IL6 (Servicebio, GB11117, 1:500), and rabbit anti-IL10 (Servicebio, GB11108, 1:200). After washing with PBS (Servicebio, G0002), slices were stained with donkey anti-rabbit IgG (Servicebio, GB21403, 1:300), goat anti-rabbit IgG (Servicebio, GN25303, 1:400), and goat anti-rabbit (Servicebio, GB23303, 1:200) secondary antibodies for 50 min at room temperature.

After washing again with PBS (Servicebio, G0002), DAPI solution (Servicebio, G1012) was added for 10 min at room temperature to stain cell nuclei. After washing with PBS (Servicebio, G0002), autofluorescence quenching reagents (Servicebio, G1221) was added for 5 min.

### Statistical Analysis

The data are presented as the means ± standard deviations. Statistical analysis was conducted via one-way or two-way analysis of variance (ANOVA) via GraphPad Prism 10 and Origin 2025. p > 0.05 was considered not significantly different (ns, not significant; *p < 0.05, **p < 0.01, and ***p < 0.001).

## Acknowledgements

We gratefully acknowledge the support of Novo Nordisk Foundation (NNF22OC0080508), Lundbeck Foundation (R400-2022-1232), Carlsberg Foundation (CF19–0300), Marie Skłodowska-Curie grant agreement ENSIGN (101086226), NanoRam (101120146) and L4DNANO (101086227) for the support.

