## supplementary materials for "Hydrogel–Electromagnetic Biohybrid Systems Direct Neural Morphogenesis across Central and Peripheral Systems" (from another version of this preprint: version 3, 2025-11-01)

**Supplementary information**

**Oriented Hydrogels with Wireless Electromagnetic Cues Direct Neural Morphogenesis Across Central and Peripheral Systems**

Bjarke Nørrehvedde Jensen^1^, Tong Tong^2,3^, Yun Tang,^4^ Grith Skovborg^1^, Yuge Zhang,^5^ Kjeld Kaj Klompmaker^1^, Cecillie Linneberg Matthiesen^1^, Jens Toft Eschen^2,3^, Kirstine Juul Elbæk^2,3^, Yuting Wang^1^, Lone Tjener Pallesen^6^, Dror Seliktar^7^, Di Sun,^4^ Mingdong Dong^5^, Christian Bjerggaard Vægter^6^, Anders Rosendal Korshøj^2,3^, Menglin Chen^1,5^*

*^1^ Department of Biological and Chemical Engineering, Aarhus University, Aarhus, Denmark*

*^2^ Department of Neurosurgery, Aarhus University Hospital, Palle Juul-Jensens Boulevard 165, 11 8200 Aarhus N, Denmark*

*^3^ Department of Clinical Medicine, Aarhus University, Palle Juul-Jensens Boulevard 99, 8200 13 Aarhus N, Denmark*

*^4^ Department of Ultrasound in Medicine, Shanghai Institute of Ultrasound in Medicine, Shanghai Key Laboratory of Neuro-Ultrasound for Diagnosis and Treatment, Shanghai Sixth People’s Hospital Affiliated to Shanghai Jiao Tong University, School of Medicine, Shanghai Jiao Tong University, Shanghai, China*

*^5^ Interdisciplinary Nanoscience Center, Aarhus University, Aarhus, Denmark*

*^6^Department of Biomedicine, Aarhus University, Aarhus C, Denmark*

*^7^ Faculty of Biomedical Engineering, Technion – Israel Institute of Technology, Haifa 32000, Israel*

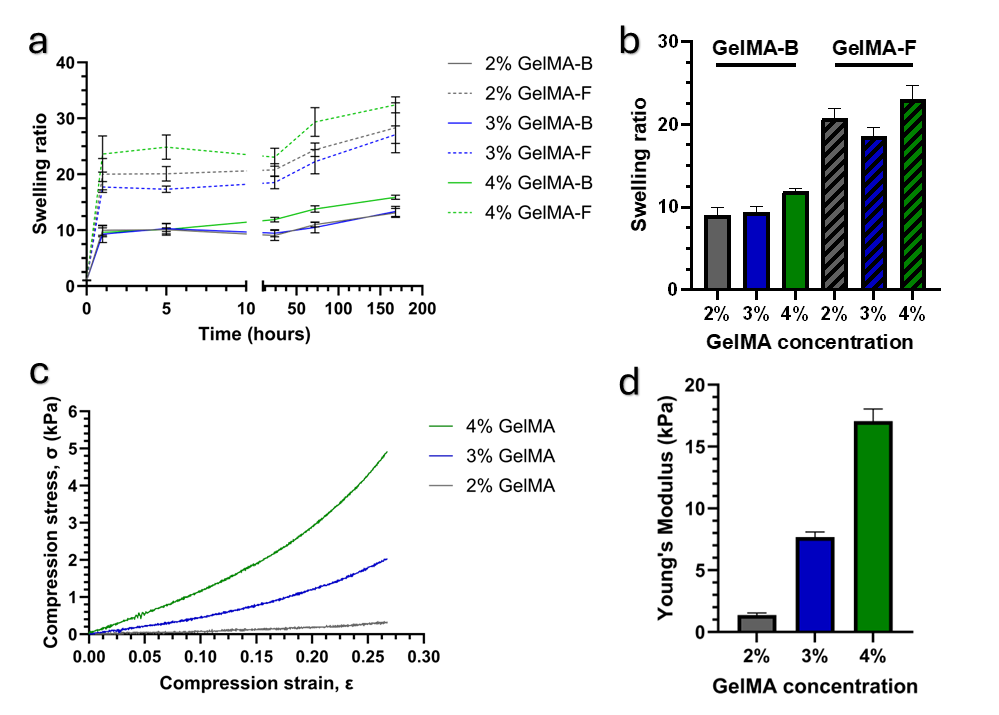

**Figure S1 – Rheological measurements of GelMA. a*)*** *Swelling ratios of GelMA-B and GelMA-F over time in PBS.* ***b)*** *Swelling ratios of GelMA-B and GelMA-F after 7 days in PBS.* ***c)*** Compression stress-strain curves of GelMA. ***d)*** *Young’s Modulus of GelMA from (****c****).*

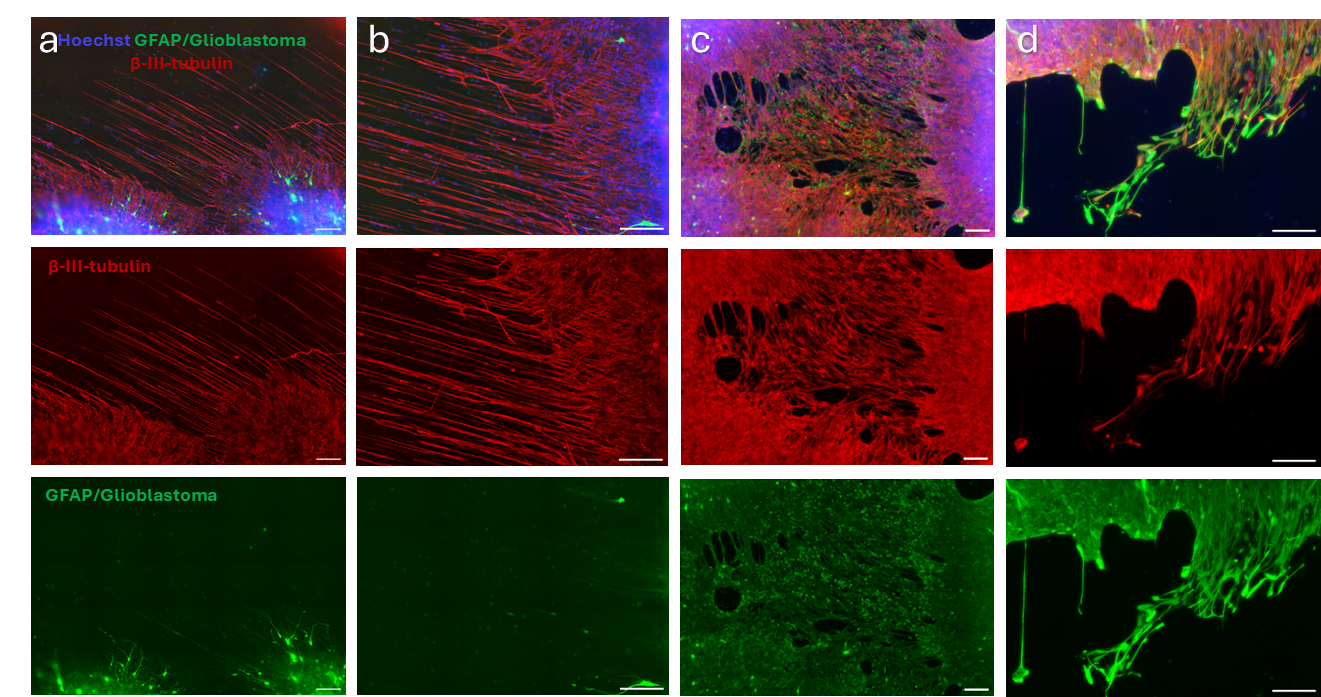

**Figure S2 – Human brain tissue on membrane well-insert and GelMA-F.** *Human brain tissue after 21 days of culture on 2% w/v* ***a-b)*** *GelMA-F and* ***c-d)*** *membrane well-insert post-stained for GFAP (green). Blue = Hoechst, red = β-III-tubulin, green = glioblastoma/GFAP. Scalebar = 200 µm.*

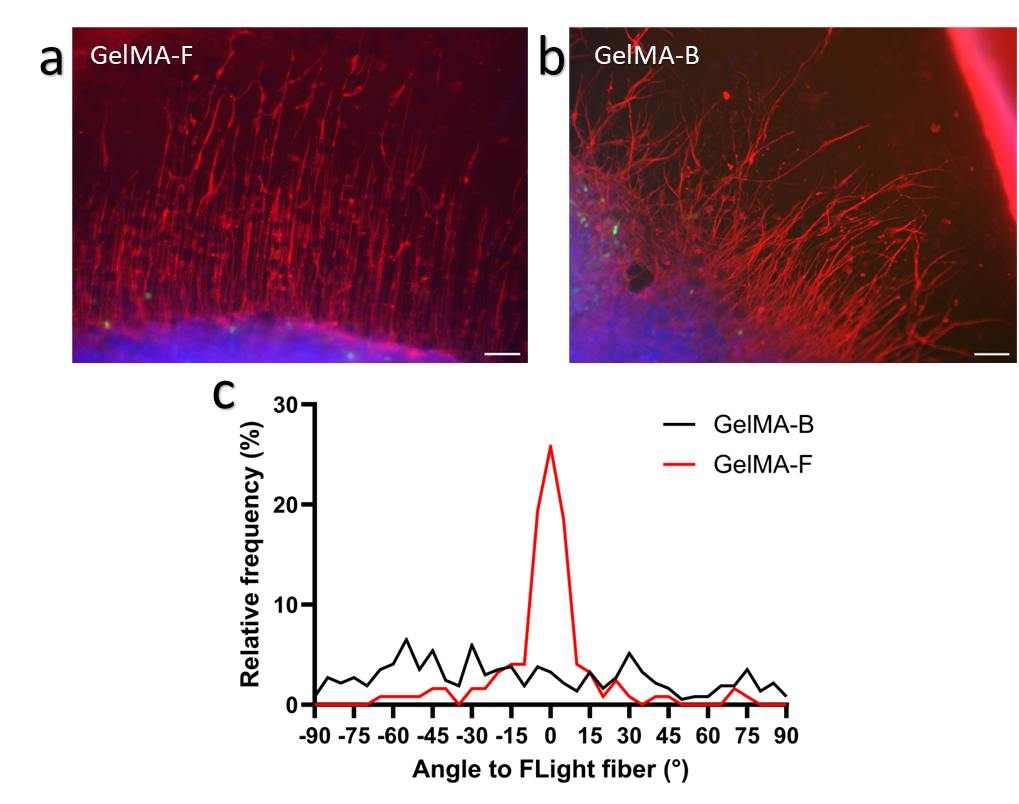

**Figure S3 – Human brain tissue on GelMA-B and GelMA-F.** *Human brain tissue after 7 days of culture on 2% w/v* ***a)*** *GelMA-F and* ***b)*** *GelMA-B. Blue = Hoechst, red = β-III-tubulin, green = glioblastoma. Scale bar = 100 µm.* ***c)*** *Outgrowth directionality analysis from* ***a)*** *and* ***b)****.*

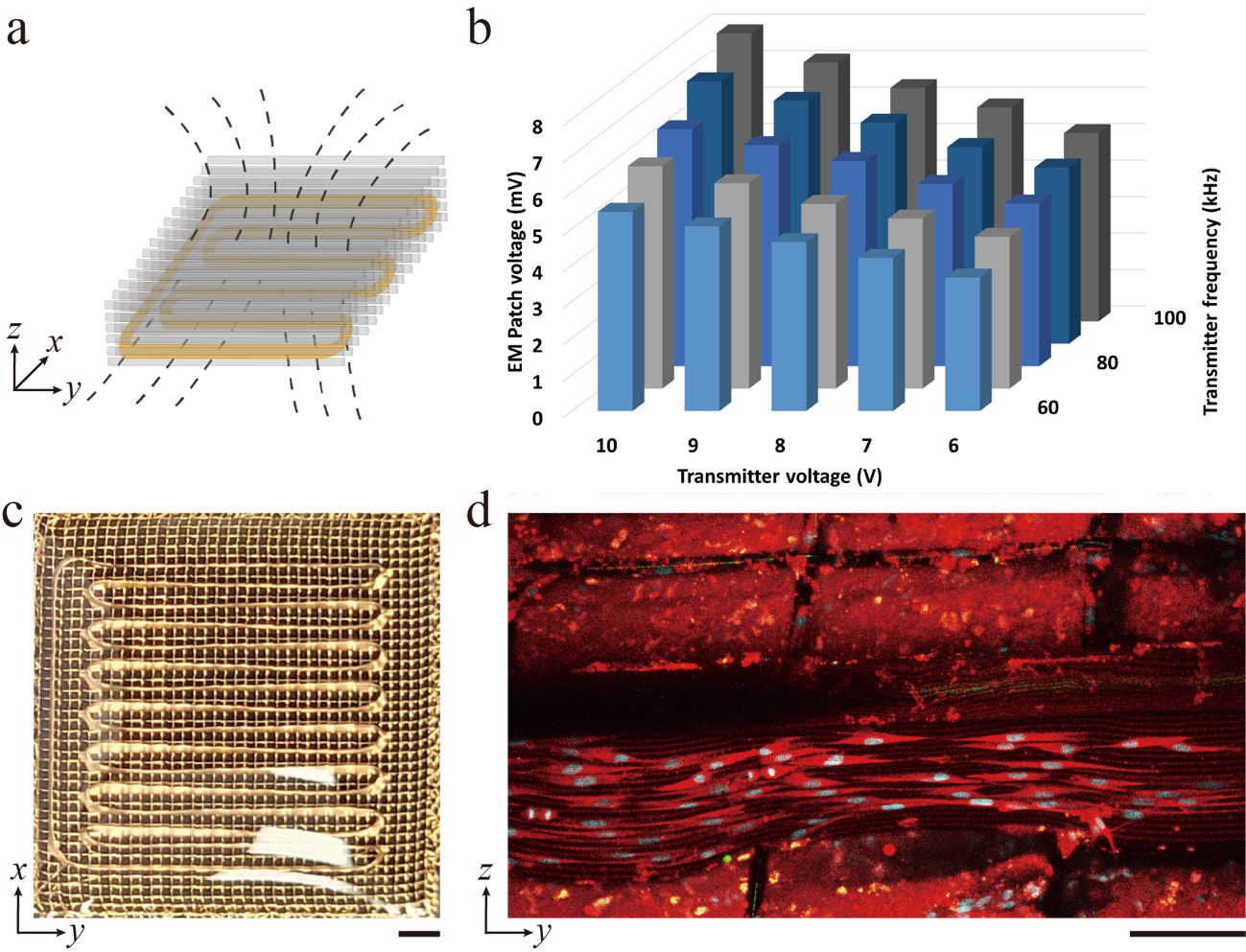

**Figure S4 *a)*** *Schematic of EM patch.* ***(b)*** *Voltage output of EM patch encapsulated in GelMA.* ***(c)*** *Photograph of hydrogel encapsulated EM patch.* ***(d)*** *ICC fluorescence image of cell migration from brain tissue onto stimulation interface of the EM patch, blue = Hoechst, red = β-III-tubulin, green = glioblastoma/GFAP, Scalebar = 100 µm.*

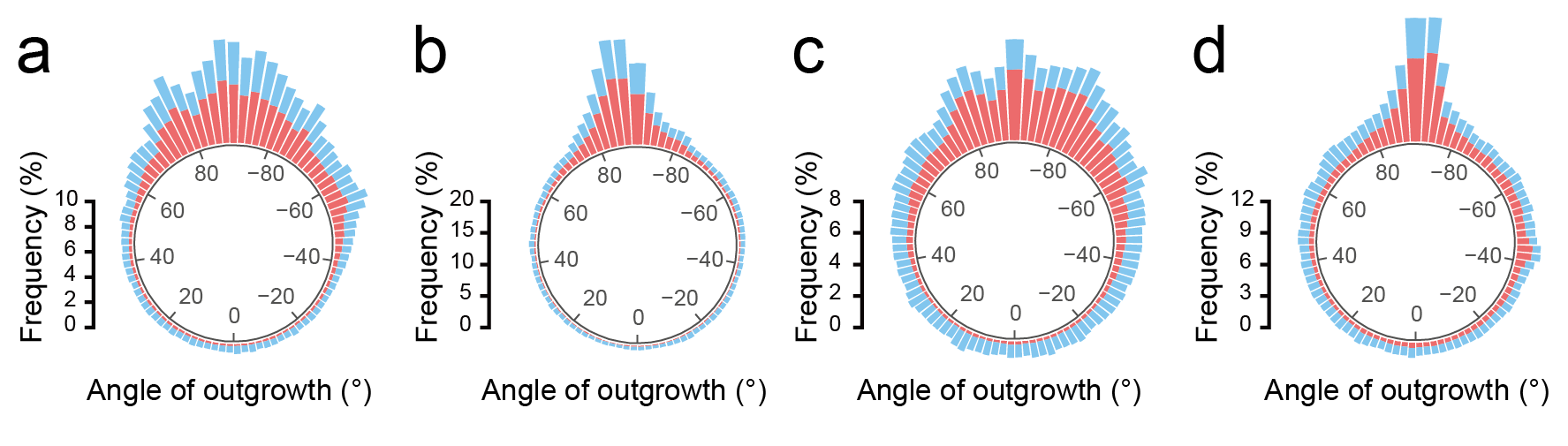

**Figure S5** *DRG explants outgrowth directionality analysis of GelMA-B (a), GelMA-F (b), GelMA-B stimulation (c) and GelMA-F stimulation (d).*

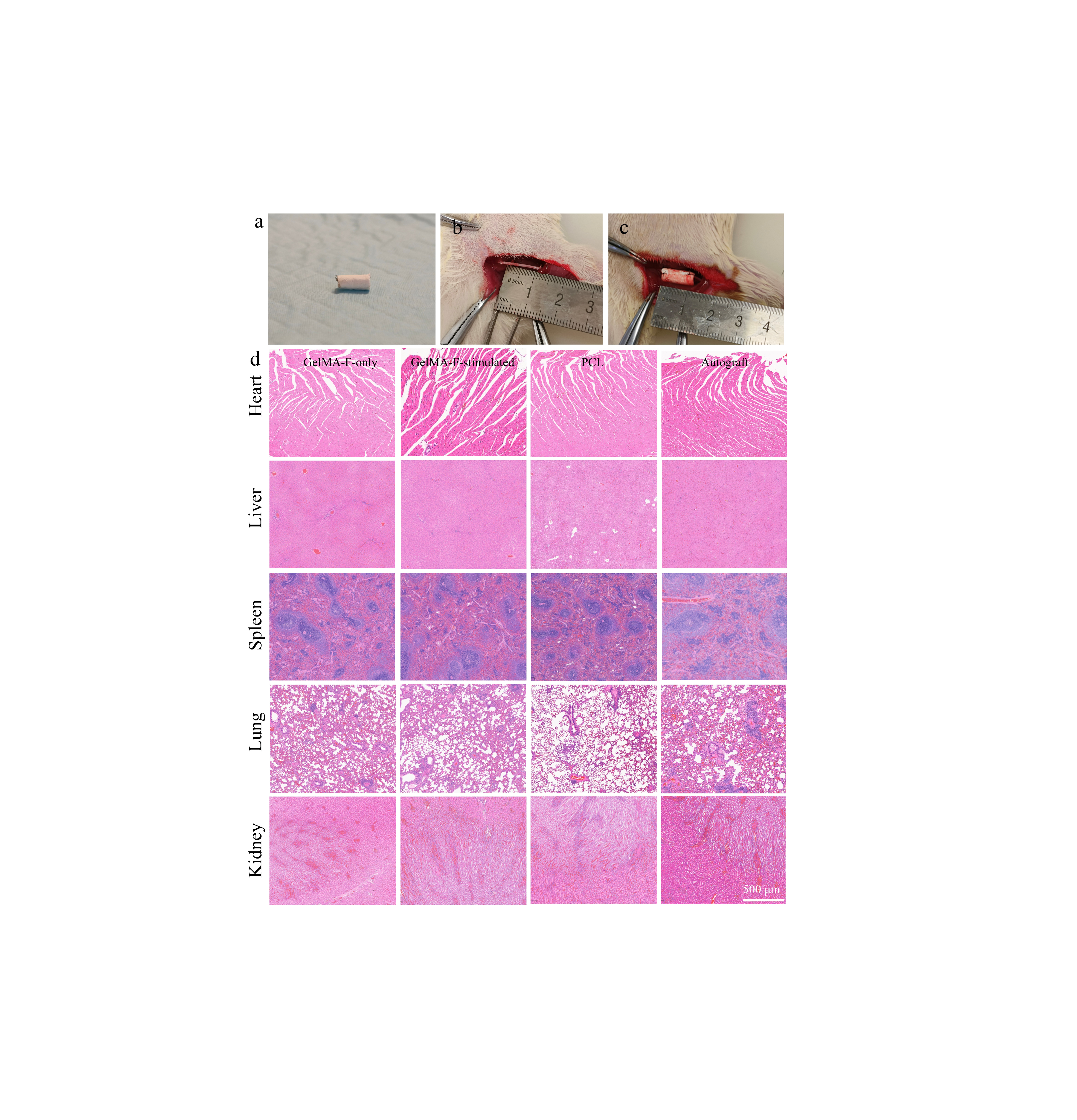

**Figure S6.** Surgical implantation and systemic biocompatibility evaluation in a rat sciatic nerve defect model. (**a**) Representative image of the fabricated GelMA-F nerve guidance conduit. (**b, c**) Intraoperative photographs depicting the creation of a 10 mm sciatic nerve defect in a rat model (**b**) and the subsequent implantation of the nerve guidance conduit into the defect site (**c**). A surgical ruler is included for scale. (**d**) HE staining of major organs (heart, liver, spleen, lung, and kidney) collected at 8 weeks post-implantation from all experimental groups. All organ sections consistently exhibit normal tissue architecture, indicating no apparent systemic toxicity or inflammatory responses associated with the respective treatments. Scalebar in (**d**) = 500 µm.

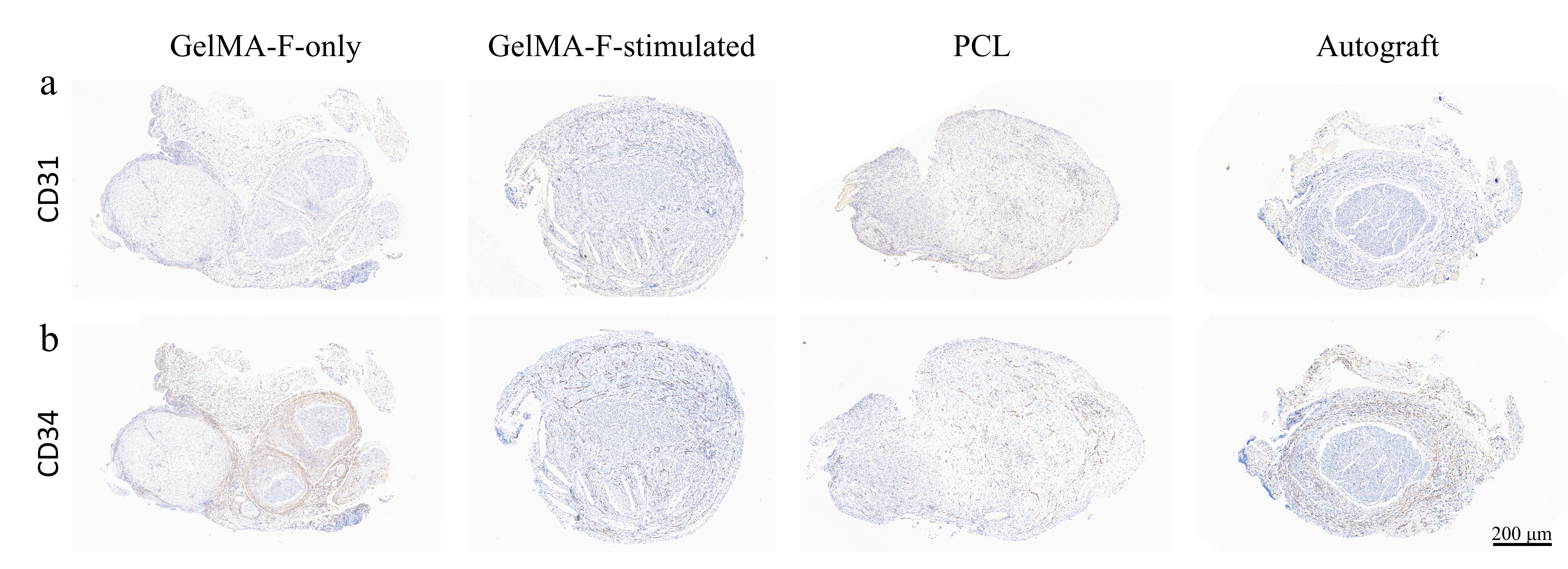

**Figure S7.** Representative immunohistochemical staining of regenerated sciatic nerve and images display positive staining (brown) for: (**a**) CD31 and (**b**) CD34, indicative of nascent vasculature. Sections were counterstained with hematoxylin (blue) to visualize cell nuclei. Experimental groups, including GelMA-F-only, GelMA-F-stimulated, PCL, and Autograft, are indicated above the respective columns. Scalebar = 200 µm.

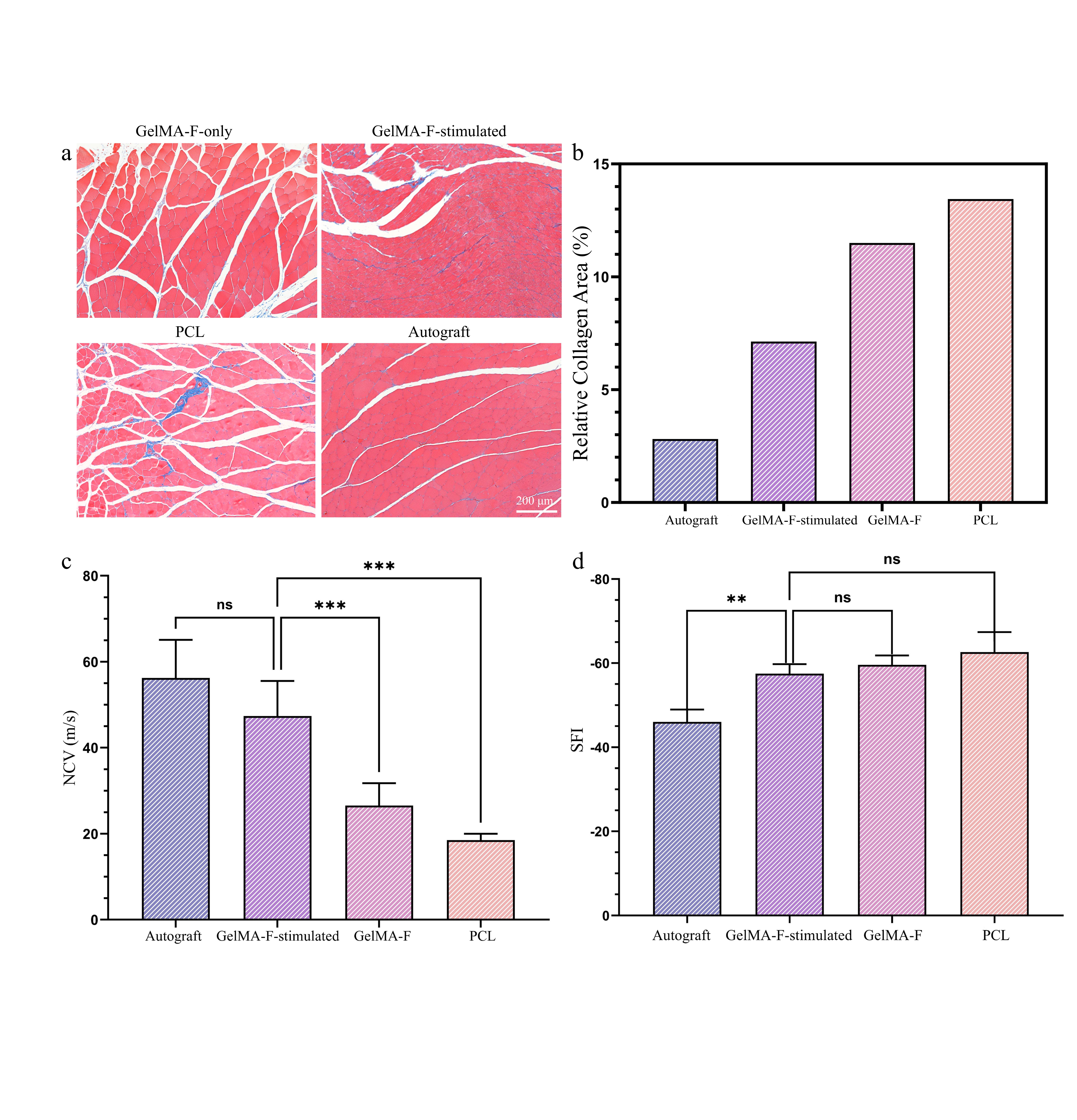

**Figure S8.** Histological assessment of gastrocnemius muscle regeneration and functional recovery. (**a**) Representative Masson’s Trichrome staining of transverse gastrocnemius muscle sections, and muscle fibers are stained red/pink, while collagen deposition, indicative of fibrosis, appears blue. (**b**) Quantification of the relative collagen area (%) within the gastrocnemius muscle sections.

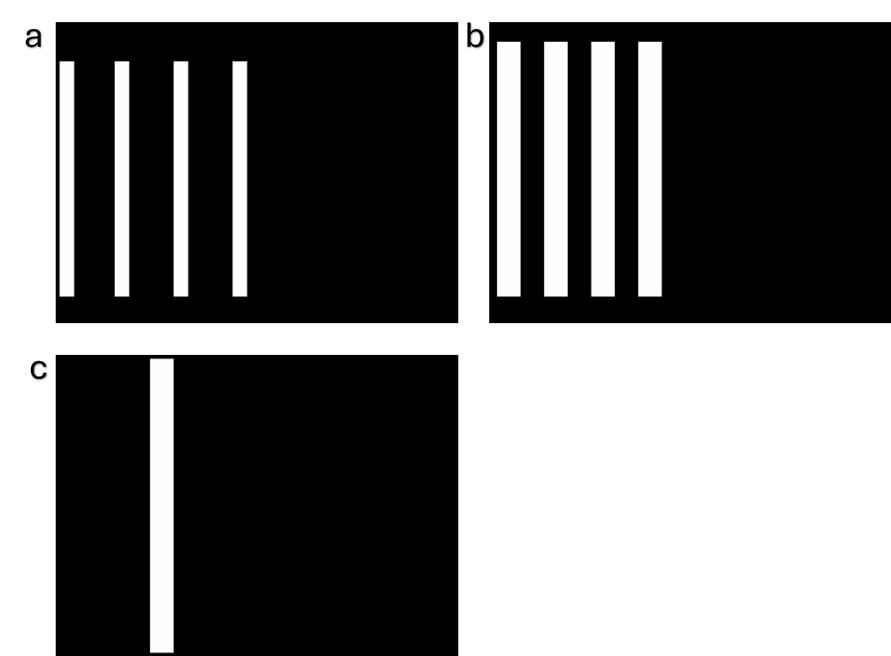

**Figure S9 – Printing exposure files for GelMA-F fabrication.** ***a)*** *1000 µm thick slices used for human brain slice culturing.* ***b)*** *1400 µm thick slices used for encapsulation of eGFP-MSC and for seeding dissociated DRG neurons onto.* ***c)*** *1400 µm thick slice used for encapsulation of EM stimulation patch for seeding DRGs onto for wireless EM stimulation.*

**Table S1 – G-code for melt electrowriting of the EM coil.**

#1=750

#2=50

#3=50

G90

F#1 G1 X0 Y0

G91

F#1 G1 X1

M98 P06 L1

F#1 G1 X-1

M30

O06

M98 P07 L2

F#1 G1 X1 Y1.5

M98 P01 L30

F#1 G1 X-1 Y-1.5

M99

O07

M98 P08 L20

M98 P09 L20

F#2 G1 Y-10

F#2 G1 X-10

F#2 G1 Y10

F#2 G1 X10

F#1 G1 Y-10

F#1 G1 X-10

M99

O08

G91

F#1 G1 X10

F#3 G1 Y0.25

F#1 G1 X-10

F#3 G1 Y0.25

M99

O09

G91

F#1 G1 y-10

F#3 G1 x0.25

F#1 G1 y10

F#3 G1 x0.25

M99

O01

G91

M98 P02 L8

F#1 G1 Y-0.5

F#1 G1 X-8

F#1 G1 Y0.5

M99

O02

G91

F#1 G1 Y8

F#1 G1 X0.5

F#1 G1 Y-8

F#1 G1 X0.5

M99

O40

G91

F#1 G1 X40

F#1 G1 X-40

M99
